# Spatial transcriptomics reveals epithelial–immune remodeling preceding malignant transformation of oral premalignant lesions

**DOI:** 10.64898/2026.09.17.751597

**Authors:** Yuxiao Jarvan Jiang, Irshad Ali, Jeanine M. Genkinger, Oliver D. King, Elizabeth Phillipone, Fatemeh Momen-Heravi

## Abstract

Oral premalignant lesions (OPLs) are common; however, histopathological grading incompletely identifies lesions destined for cancer. To define tissue ecosystems that precede malignant transformation, we integrated single-cell-resolution Xenium spatial transcriptomics of 20 HPV-negative biospecimens from 16 patients, with independent single-cell RNA sequencing and immunofluorescence data from 38 patients with OPLs after quality control. Progressive OPLs exhibit a coordinated epithelial–immune program comprising MX1/NOTCH3-high KRT14⁺/KRT15⁺ basal epithelial states, S100A9-high inflammatory macrophages, TIGIT-high, exhaustion-associated T cells, and altered dendritic cell states. Spatial analyses showed that T cells and dendritic cells were displaced from the basal epithelial interface, and that epithelial neighborhoods were depleted of dendritic cells but enriched for TIGIT-high T cells, revealing immune reorganization not captured by cellular abundance alone. Independent single-cell and protein-level analyses reproduced basal epithelial interferon/stress programs, S100A9-associated myeloid inflammation, TIGIT-high, exhaustion-associated T-cell programs, and dendritic cell redistribution. Ligand–receptor inference suggested convergent myeloid and lymphoid signals linked to epithelial stress, checkpoint regulation, and extracellular matrix remodeling. Functionally, recombinant S100A9 accelerated wound closure in oral epithelial and cancer cells, whereas S100A9 and S100A8/S100A9 enhanced oral cancer cell proliferation. These responses were attenuated by pharmacologic inhibition of TLR4 or RAGE. Together, these data define a spatially organized, myeloid-skewed, checkpoint-enriched ecosystem present before invasion and suggest S100A9–TLR4/RAGE signaling and epithelial–immune geometry as candidate mechanisms and biomarkers for oral cancer interception.

**Ethical Compliance:** All procedures performed in studies involving human participants were in accordance with the ethical standards of the institutional and/or national research committee and with the 1964 Helsinki Declaration and its later amendments or comparable ethical standards.

**Statement of significance:** Oral premalignant lesions are common, but histopathologic grading cannot reliably identify which will become cancer. By combining spatial and single-cell transcriptomics of lesions with known clinical outcomes and validating the findings in independent cohorts, we identify a preinvasive ecosystem that precedes malignant transformation. This ecosystem includes stressed basal epithelial cells, S100A9-high inflammatory macrophages, TIGIT-high, exhaustion-associated T cells, and displacement of dendritic cells and T cells from the epithelial interface. In functional assays, S100A9 enhanced epithelial-cell migration and proliferation, and these responses were attenuated by pharmacologic inhibition of TLR4 or RAGE. These findings show how epithelial stress, inflammation, and impaired immune surveillance converge before invasion and may support improved risk assessment and strategies to prevent oral cancer.

## INTRODUCTION

Oral squamous cell carcinoma (OSCC) is one of the most common types of head and neck squamous cell carcinoma (HNSCC) and remains a major global health burden (*1*). The prognosis of OSCC still depends on the stage at diagnosis, underscoring the need for earlier biology-informed detection strategies (*1*). Many human papillomavirus (HPV)-negative OSCCs arise from oral premalignant lesions (OPLs); however, current risk stratification relies heavily on histopathology, which incompletely captures the molecular ecology that drives progression (*2*).

Large-scale tumor profiling has established the mutational and pathway architecture of HNSCC, including tobacco-associated *TP53*-mutant disease with recurrent *CDKN2A*, *NOTCH*, and *PI3K* alterations, as well as oxidative-stress programs that likely emerge already in the premalignant continuum (*3*). At single-cell resolution, *in vivo* atlases of HNSCC have revealed epithelial heterogeneity, immune cell states, and stromal programs that together shape tumor behavior (*4*).

Spatially resolved transcriptomics provides a powerful approach for mapping cell states in situ while preserving tissue architecture (*5, 6*). When paired with single-cell RNA sequencing (scRNA-seq) and ligand–receptor inference frameworks, spatial profiling can identify candidate cell–cell communication programs and immune– epithelial interactions that may precede malignant transformation (*7*). In the oral mucosa, epithelial compartments, stromal subsets, and immune circuits form specialized barrier niches that regulate tissue immunity and disease-related cell states (*8*). In parallel, studies of cancer evolution and oral carcinogenesis suggest that immune selection, immune checkpoint expression, and T-cell dysfunction can emerge before frank invasion, supporting a model in which immune microenvironments shape premalignant evolutionary trajectories (*9–11*).

At the clinical interface, histologic grading of OPLs remains variably reproducible and only moderately predictive; inter-observer agreement is limited even among experienced oral pathologists, particularly across intermediate grades (*12, 13*). Contemporary reviews and meta-analyses indicate that transformation risk reflects a composite of histologic grade, clinical phenotype, anatomic site, and tobacco/alcohol exposure; however, current models remain insufficient for confident patient-level decisions (*13, 14*). These limitations have motivated prognostic frameworks that integrate quantitative, biology-anchored features, including molecular biomarkers, immune cell states, image-derived morphometrics, and spatial tissue context. Thus, the unresolved biological gap is not only the epithelial or immune cells present in OPLs but also how dysplastic epithelium and immune surveillance are spatially organized before invasion to differentiate which OPLs will progress to OSCC.

Here, we built an FFPE-compatible multimodal atlas of the OPL-to-OSCC continuum by integrating Xenium spatial transcriptomics with independent scRNA-seq and immunofluorescence validation cohorts. We profiled non-progressive oral premalignant lesions (NP-OPL; no malignant transformation over ≥5 years of follow-up), progressive oral premalignant lesions (P-OPL; malignant transformation to OSCC within 5 years of the index biopsy), and OSCC biopsies to define epithelial and immune cell states associated with progression, quantify epithelial-neighborhood composition and epithelial–immune distances, and identify candidate ligand–receptor axes that may mediate immune dysfunction and epithelial remodeling. By focusing on progression-associated programs that are reproducible across spatial, single-cell, and protein-level validation, this study identifies early epithelial–immune niche remodeling as a measurable feature of malignancy risk and provides a foundation for biomarker-guided surveillance and precision interception in oral cancer.

## METHODS

### Sample Acquisition

Formalin-Fixed Paraffin-Embedded (FFPE) blocks of biopsies from patients with at least 5 years of follow-up with a histological diagnosis of non-progressive oral premalignant lesions (NP-OPLs), progressive oral premalignant lesions (P-OPLs), or oral squamous cell carcinoma (OSCC) were acquired from 55 patients based on the exclusion and inclusion criteria. NP-OPLs were defined as lesions with no malignant transformation over a minimum clinical follow-up of 5 years, whereas P-OPLs were defined as lesions that progressed to OSCC within 5 years of index biopsy. Patients with histologically confirmed oral premalignant lesions and available FFPE tissue of sufficient quality for spatial transcriptomic, scRNA-seq, or immunofluorescence analysis were considered eligible. The exclusion criteria included insufficient tissue, poor fixation or section quality, failed assay QC, inadequate clinical follow-up for NP-OPL classification, HPV-positive lesions, ambiguous progression status, prior invasive OSCC at the same site before the index premalignant biopsy and missing essential clinical annotation.

Disease status (NP-OPL, P-OPL, and OSCC) and demographic and clinical information, including age at biopsy collection, progression date (when applicable), sex, ethnicity, race, histological findings, time to malignant progression, recurrence, OSCC diagnosis (for OSCC samples), biopsy site, and history of tobacco and alcohol consumption, were retrieved from the electronic health record. All studies were approved by the IRB of Columbia University (protocol# AAAS0169).

### Study design, cohorts, and reporting structure

This study analyzed FFPE oral lesion biospecimens across three assay arms: Xenium spatial transcriptomics (discovery), single-cell RNA sequencing (validation), and immunofluorescence (validation). The Xenium discovery cohort included eight NP-OPL biospecimens, eight P-OPL biospecimens collected before malignant transformation, and four subsequent OSCC biospecimens longitudinally matched at the patient level to four of the eight P-OPL biospecimens. Thus, four patients contributed paired P-OPL and OSCC biospecimens, whereas the remaining biospecimens were contributed by unique patients. The independent immunofluorescence validation cohort included 23 patients and biospecimens, comprising 12 NP-OPL and 11 P-OPLs. The lesion sites were abstracted from pathology reports and surgical notes, and the histologic grade was reviewed and recorded by an oral pathologist. All the included samples were HPV-negative. Cohort membership, clinical and demographic variables, lesion site, histologic grade, and history of alcohol consumption and smoking are provided in Table S1. No statistically significant differences in age, biological sex, alcohol or smoking status categories (former, current, missing), or lesion site (high-risk versus low-risk) were detected between the cohorts of P-OPL and NP-OPL. Age was compared using the two-tailed Mann–Whitney U test for two-group comparisons. Categorical variables, including biological sex and lesion site, were compared using Fisher’s exact test or Fisher– Freeman–Halton exact test for multi-category contingency tables.

### Xenium in situ workflow

Xenium in situ gene expression assays use premade gene panels. DNA probes for target genes contain two regions complementary to the target RNA flanking a gene-specific barcode, and are circularizable for circular amplification. In this study, we used a pre-designed Xenium Human Immuno-Oncology Profiling Panel with 380 target genes. FFPE slide preparation followed stepwise instructions in the 10x Genomics Demonstrated Protocol for FFPE Tissue Preparation Guide CG000578. FFPE blocks were rehydrated and sectioned with a microtome into 5µm sections on Xenium slides. Three to four sections were placed on each Xenium slide. The slides were dried overnight before deparaffinization and decrosslinking according to the 10x Genomics Demonstrated Protocol CG000580. Probes of the immuno-oncology panel were then hybridized to the slides using the Probe Hybridization Mix by incubating at 50 °C overnight on a thermal cycler. Following hybridization, the slides were washed with PBS-T and incubated at 37 °C with post-hybridization buffer for 30 min before ligation and amplification. After amplification, the slides were washed with PBS and processed for autofluorescence quenching and nuclear staining. 10x Genomics Demonstrated Protocol CG000582 provided the buffer components and stepwise procedures for probe hybridization, ligation, and amplification.

#### Xenium Analyzer

Xenium Analyzer onboard processing was performed according to the 10x Genomics Demonstrated Protocol CG000584. To perform spatially resolved transcriptomic profiling on the Xenium Analyzer, FFPE tissue sections were mounted on Xenium slides, placed in cassette wells containing 1 mL PBS-T, and loaded onto the instrument, where an overview scan was acquired to delineate the regions of interest across a grid of fields of view. The system then executed iterative cycles of fluorescent probe hybridization, high-resolution imaging, and probe removal, capturing vertical image stacks for every channel and field that were automatically flat-field-corrected, stitched, and registered on-board. Throughout the run, puncta corresponding to individual transcripts were detected and decoded in real time to assign gene identities and quality scores. Upon completion, the instrument performed an automated fluidics cleanup and exported a comprehensive data bundle containing per-transcript spatial coordinates, cell-by-gene count matrices, and quality metrics for visualization in Xenium Explorer. Cell segmentation was performed using a Xenium Analyzer based on DAPI-stained nuclei. Cell boundaries were defined by expanding the boundaries of the stained nuclei by 15µm or until contact with other cell boundaries. After the Xenium in situ assays, the slides were processed with post-Xenium H&E staining according to the stepwise guidelines from the 10x Genomics Demonstrated Protocol CG000613. The slides were stored in PBS-T at 4 °C for less than 1 week before H&E staining.

#### Data Pre-processing

Post-Xenium H&E staining images were overlaid with Xenium DAPI-stained morphology images in Xenium Explorer 3. Approximately 100 anchors were set on cell landmarks with discernible features for paired H&E and Xenium images from each region of interest. Xenium Explorer 3 automatically aligned the H&E and Xenium images according to these anchors.

### scRNA analysis pipeline

#### Data Loading and Quality Filtering

For each biospecimen, the Xenium in situ assay generated 10x-format output files (barcodes, features, and matrix) suitable for downstream analysis with single-cell RNA sequencing pipelines and a Xenium output file for analysis with the Xenium spatial analysis pipeline. The 10x-format files from each biospecimen were loaded into Seurat v5 and converted into Seurat objects labeled with a unique biospecimen identifier and disease status (NP-OPL, P-OPL, or OSCC). Initial quality filtering was performed to retain genes expressed in at least three cells and cells expressing 20–80 genes with fewer than 450 total counts. After QC, 20 of the 20 biospecimens were retained for analysis. The final Xenium dataset contained 523,753 cells, 41,326,882 decoded transcripts, and 380 genes from the 380-gene panel, with a median of 55 transcripts per cell and 30 genes per cell. The per-biospecimen QC metrics, including raw and retained cell numbers, transcript counts, median genes per cell, median transcripts per cell, and exclusion reasons, are provided in Table S2. Each Seurat object that passed the quality filtering was pre-processed using the NormalizeData (with normalization method “LogNormalize”) and FindVariableFeatures (with selection.method “vst”) functions. Variable genes used for integrating the Seurat object were identified using the SelectIntegrationFeatures function, and ScaleData and RunPCA functions were applied to each object using these genes. The resulting Seurat objects were then integrated into a single Seurat object using the “rpca” method.

#### Dimension Reduction, Clustering, and Cell Type Annotation

Seurat v5 was used to perform dimension reduction, clustering, and cell type annotation of the integrated Seurat object. The integrated Seurat object was normalized by “LogNormalize.” Dimension reduction was performed using PCA, and the first 15 principal components were retained for downstream analysis based on the elbow plot bend point. Cells in the dimension-reduced PCA space were clustered using the Louvain algorithm and visualized using a Uniform Manifold Approximation and Projection (UMAP) plot. Each cluster was annotated by matching its top differentially expressed genes, identified using FindAllMarkers (Wilcoxon rank-sum test, one-vs-rest), to canonical signature genes for epithelial cells (*EGFR*, *EPCAM*, *ERBB2/3*, *CEACAM*s), fibroblasts (*FN1*), endothelial cells (*PLVAP*, *SPARCL1*), T cells (*IL2RB*, *TRAT1*), macrophages (*MPEG1*, *CD163*), plasma cells (immunoglobulin), mast cells (*CPA3*, *P2RX1*), B cells (*MS4A1*, *PAX5*), dendritic cells (*CD1A*, *CD1C*), and vascular cells (*RGS5*).

#### Subclustering and Subcluster Annotation

Clusters identified by the Louvain algorithm were subsetted from the integrated Seurat object according to their cell type and were further sub-clustered using the abovementioned strategy. Each subcluster was characterized by its top differentially expressed markers (FindAllMarkers, Wilcoxon rank-sum test, one-vs-rest), defined as genes ranked among the top five by positive log2FC with an adjusted p-value < 0.1. These top-5 marker sets did not overlap across the subclusters. For each biospecimen, the proportion of cells belonging to each cluster and subcluster was tabulated and exported as a CSV file for downstream statistical analysis.

### Spatial Analysis Pipeline

#### Data Loading and Quality Filtering

Xenium output files from each biospecimen were loaded into Seurat v5 by using the “LoadXenium” function. Each Seurat object was labeled with a unique biospecimen identifier and disease status (NP-OPL, P-OPL, or OSCC), and the cell IDs were prefixed with the corresponding biospecimen identifier. Seurat objects were filtered to retain only cells with gene counts between 10 and 100 and total transcript counts less than 1000. The filtered Seurat objects were merged into a single Seurat object.

#### Dimension Reduction, Clustering, and Cell Type Annotation

Counts in the merged Seurat object were normalized and scaled using SCTransform. Dimension reduction was performed using PCA, and the number of principal components was selected from an elbow plot generated across the first 20 PCs; 15 PCs were retained for downstream analysis. Cells were clustered in PCA space, and UMAP was used to visualize the resulting clusters. The merged Seurat object was processed using PrepSCTFindMarkers, and cluster marker genes were identified using FindAllMarkers. Clusters were annotated using canonical markers for epithelial cells (*EGFR, ERBB2/3*), fibroblasts (*FN1, PDGFRA, SPARC*), and endothelial cells (*PLVAP, SPARCL1*), together with spatial localization in Xenium images. Immune cells were annotated using SingleR with the Monaco immune reference (*15*) and confirmed using top markers for T cells (*CD3D, CD3E*), macrophages (*MPEG1, CD163*), neutrophils (*CXCL1*), plasma cells (*MZB1* and immunoglobulin genes), Natural-Killer (NK) cells (*KLRF1, KLRC1, GNLY*), mast cells (*CPA3, MS4A2*), B cells (*MS4A1, CD79A*), and dendritic cells (DCs) (*FCER1A, CLEC10A*). Clusters assigned to the same cell type were merged for downstream spatial analysis.

#### Pair-wise Measurement of Distance Between Cell Types of Interest

The centroid coordinates for the cell types of interest were extracted from their respective field of view (FOV) image objects in the merged Seurat object. These coordinates were converted into data frames and the Euclidean distance matrix was computed between each pair of epithelial and immune cells. The matrix quantifies the spatial proximity of each pair. The combined data frames from each biospecimen were further consolidated by disease group and filtered to retain only cell pairs with a distance of less than 100 µm. Histogram plots were generated, and a statistical summary, including mean, median, minimum, maximum, and percentiles, was exported for data presentation.

#### Nearest Neighbors Analysis

For each field of view, epithelial cells were defined as the target population and immune cells as potential neighbors. For each epithelial cell, the k-nearest immune neighbors were identified from spatial coordinates and retained if within 100 µm; k = 5 was used to characterize the overall immune neighborhood, and k = 2 to identify the most proximal T-cell contacts. Neighbor composition was quantified as raw counts and relative percentages, and exported for downstream analysis.

### Whole-transcriptome Single-Cell RNA Sequencing Validation

The independent validation cohort consisted of 16 FFPE oral premalignant lesion biospecimens from 16 unique patients, including 11 P-OPL biospecimens and 5 NP-OPL biospecimens **(Table S1)**. Each patient contributed only one specimen; therefore, the validation cohort contained no paired samples or repeated measures. The validation cohort was analyzed separately from the Xenium spatial transcriptomics discovery cohort and used to evaluate whether progression-associated markers identified in the spatial dataset were reproducible using whole-transcriptome single-cell RNA sequencing.

#### Sample Processing

For each specimen, 2–3 scrolls of 50-µm FFPE tissue sections, depending on the available tissue quantity, were prepared from FFPE-preserved oral dysplastic lesion samples and subjected to single-cell RNA sequencing using the 10x Genomics Chromium Single Cell Gene Expression Flex protocol. Sequencing was performed with a target recovery of approximately 8,000 cells per sample and approximately 15,000 reads per cell.

#### Data Loading

10x format files of each sample were loaded into Seurat v5 and converted into Seurat objects with object names based on sample index. The initial quality filtering of each object was performed to retain genes expressed in at least three cells and cells expressing at least 200 genes. Objects with the same disease status were merged into a large Seurat object. The two large Seurat objects (i.e., P-OPL and NP-OPL) were filtered to retain cells with fewer than 5000 genes, fewer than 10000 counts, and less than 5% mitochondrial RNA; they were then pre-processed with log normalization and FindVariableFeatures. After QC filtering, the scRNA-seq validation dataset retained 54,096 cells from 5 NP-OPL and 10 P-OPL biospecimens (one of the 11 P-OPL samples was excluded for failing quality control), including 21,313 epithelial cells, 18,523 immune cells, 6,963 fibroblasts, 4,595 endothelial cells, and other minor cell types. The per-sample cell numbers, sequencing depth, median genes per cell, median UMIs/counts per cell, and mitochondrial percentage are provided in Table S3. The two disease-status objects (NP-OPL and P-OPL) were normalized (LogNormalize), and their top 4,000 variable features were identified (vst). Integration anchors between the two objects were identified using canonical correlation analysis (CCA; FindIntegrationAnchors), and the objects were combined into a single integrated Seurat object using IntegrateData (LogNormalize, dims = 1:50). Downstream dimension reduction, clustering, and cell-type annotation were performed on the integrated object.

#### Dimension Reduction, Clustering, and Cell Type Annotation

Dimension reduction was performed on the integrated Seurat object by PCA using the first 50 principal components, consistent with integration dimensionality. Cells were clustered using the Louvain algorithm and visualized using UMAP. Each cluster was annotated with a general cell type based on signature genes for epithelial cells (*KRT5*, *KRT14*, *S100A2*, *CSTA*, *SPRR1B*), endothelial cells (*ACKR1*, *RAMP2*, *SELE*, *VWF*, *PECAM1*), fibroblasts (*LUM*, *COL3A1*, *DCN*, *COL1A1*, *CFD*), immune cells (*CD69*, *CD52*, *CXCR4*, *PTPRC*, *HCST*), as reported in a previous study on oral tissue (*8*), and also muscle cells (*MYH1*, *MYL1*, *MYPN*, *MYOG*, *MYLPF*), salivary gland cells (*ZG16B*, *PIGR*, *MUC7*, *LTF*, *TCN1*), and melanocytes (*TYR*, *TYRP1*, *MLANA*, *DCT*, *PMEL*). Cluster identities were assigned by matching these canonical signature genes to the top markers identified using FindAllMarkers (Wilcoxon rank-sum test, one-vs-rest).

#### Subclustering of Immune Cells and Immune Cell Type Annotation

Cells annotated as immune cells in the global clustering above were subsetted from the integrated Seurat object and annotated at finer granularity using SingleR (*16*). The immune object was converted to a SingleCellExperiment, restricted to genes shared with the Monaco Immune Data reference (*15*), and log normalized. Each cell was labeled individually (per-cell mode; classic DE method; fine-tuning enabled) using the fine-resolution labels of the reference, which were grouped into the following immune cell types: monocytes/macrophages, myeloid dendritic cells (mDC), plasmacytoid dendritic cells (pDC), neutrophils, mast cells, natural killer (NK) cells, MAIT cells, γδ T cells, CD4 T cells, regulatory T cells (Tregs), CD8 T cells, B cells, and plasma cells. Signature genes for each immune cell type were subsequently identified using FindAllMarkers (Wilcoxon rank-sum test, one-vs-rest) to confirm these assignments.

#### Validation of Identified Progression Markers

Cell types of interest were subsetted from the integrated object: epithelial cells from the general cell types and macrophages and T cells from immune cells. The P-OPL markers identified in the spatial transcriptomic analysis were visualized in dot plots in the corresponding cell types to evaluate the differential expression of these markers between P-OPL and NP-OPL samples.

#### Pathway Analysis and Gene Set Enrichment Analysis

Pathway enrichment analysis for Xenium data was conducted using Enrichr (*17*) with the GO Biological Process (BP) 2026 ontology. Pathway analysis of scRNA-seq data was performed using the clusterProfiler package (*18*), and gene set enrichment analysis (GSEA) (*19*) was performed using the fgsea package (*20*) with GO BP and MSigDB Hallmark gene sets.

#### Exhaustion-Associated T-cell Transcriptional Score

To quantify an exhaustion-associated T-cell transcriptional program, we computed a module score in Seurat using AddModuleScore, based on a 17-gene exhaustion signature spanning inhibitory receptors (*PDCD1*, *CTLA4*, *HAVCR2*, *LAG3*, *TIGIT*, *CD244*, *CD160*, *BTLA*, and *VSIR*), exhaustion-associated transcription factors (*TOX*, *EOMES*, and *NR4A1*), and markers of terminal differentiation (*ENTPD1*, *CD38*, *CD101*, *CXCL13*, and *KLRG1*). Scores were computed using log-normalized data.

#### Ligand–Receptor Interaction Analysis

Immune-to-epithelial ligand–receptor signaling was inferred using NicheNet with the nichenet_seuratobj_aggregate workflow (*7*). NicheNet prior ligand–receptor, signaling, and ligand–target regulatory networks were obtained from the archived NicheNet dataset on Zenodo record 3260758 (*7*). Epithelial cells were defined as the receiver population and immune cells were defined as ligand-expressing sender populations. T cell subtypes were pooled for this analysis. The epithelial target gene set was defined as genes upregulated in epithelial cells from P-OPL compared to NP-OPL. Candidate ligands expressed by immune sender cells were ranked using NicheNet ligand activity, which estimates how well each ligand’s predicted downstream target genes explain the P-OPL-associated epithelial transcriptional program. The top 20 ranked ligands and their cognate receptors expressed by the epithelial receiver cells were retained for downstream visualization. Prioritized ligand–receptor pairs were visualized using heatmaps and alluvial diagrams, with edges of interaction weight >0.2.

### Validation in External Dataset

External validation used GSE30784 (*21*), a publicly available Gene Expression Omnibus dataset of 229 oral tissue samples spanning the disease axis: 45 histologically normal mucosa, 17 dysplasia, and 167 OSCC. Whole tissue was profiled without microdissection on Affymetrix Human Genome U133 Plus 2.0 arrays. Each sample reflects the bulk lesion rather than an isolated cell population. Normalized expression values were downloaded and used as provided.

#### Signature scoring

Discovery-derived signatures — Epi.sub.1 (*MX1, CDKN2B, NOTCH3, PTGS1, NELL2*), Mac.sub.3 (*APOE, MMP12, S100A9, ISG15, SDC1*) and a 17-gene T-cell exhaustion set (*PDCD1, CTLA4, HAVCR2, LAG3, TIGIT, CD244, CD160, BTLA, VSIR, NR4A1, TOX, EOMES, ENTPD1, CD38, CD101, CXCL13,* and *KLRG1*)— were scored per sample by single-sample gene set enrichment analysis (ssGSEA; GSVA v1.52.3) on their log₂ expression, after intersecting each signature with the features present on the platform. Enrichment scores were normalized using Z-scores across the cohort.

#### Statistical analysis

All comparisons used two-tailed Mann–Whitney tests. P values were corrected by the Benjamini–Hochberg false discovery rate (FDR) procedure across the three disease-axis contrasts within each signature or gene, and, for the marker-level comparisons to normal mucosa, across all detected exhaustion markers within dysplasia and within OSCC separately. Adjusted *p* (*q*) < 0.10 was considered significant. Effect sizes were reported as Cliff’s delta, δ = 1 − 2U/(n₁n₂), where U is the Mann-Whitney statistic and n_1_ and n_2_ are the sizes of the two disease groups being compared, with positive values of δ indicating higher expression in the later-stage group. Genes were considered below reliable detection if no probe set mapped to them, or if mean intensity or variance fell below the 25th or 10th percentile of the array-wide distribution, respectively; these were tested but shown as not determined and not included for interpretation.

### Immunofluorescence Staining

5µm thick formalin-fixed paraffin-embedded tissue sections were deparaffinized by sequential immersion in xylene (3 × 5 min), followed by rehydration through graded ethanol (100% × 2, 95%, and 80%; 5 min each), and rinsed in DPBS. Heat-mediated antigen retrieval was performed in citrate buffer by microwaving the slides at high power for three cycles of 5 min each, ensuring complete buffer coverage. The slides were allowed to cool to room temperature for at least 30 min and washed with DPBS (3 × 5 min).

Tissue sections were outlined using a hydrophobic barrier pen and blocked with 1% BSA in DPBS (approximately 300–350 μL per section) for 30 min at room temperature. Sections were incubated with a primary antibody— CD3 (Proteintech, 60181-1-Ig; 1:200), CD11c (Abcam, ab23602; 1:50), TIGIT (Proteintech, 83545-1-RR; 1:100) diluted in 1% BSA/DPBS for 1 h at room temperature, followed by overnight incubation at 4 °C in a humidified chamber. Slides were then washed in DPBS (3 × 5 min) and incubated with the secondary antibodies Alexa Fluor 488 goat anti-rabbit IgG (Invitrogen, A11008; 1:500) and Alexa Fluor 568 goat anti-mouse IgG (Invitrogen, A-11004; 1:1000) for 1 h at room temperature in the dark, followed by additional DPBS washes (3 × 5 min).

For intracellular staining, the sections were permeabilized with 0.1% Triton X-100 (Sigma-Aldrich, T9284) in DPBS for 10 min and washed (3 × 5 min). After blocking with DPBS with 1% BSA (Thermo Scientific, 37520) for 30 min, slides were incubated with a second primary antibody—MX1 (Abcam, ab207414; 1:250) or S100A9 (Abcam, ab63818; 1:200) diluted in DPBS with 1% BSA for 1 h at room temperature and overnight at 4 °C. Subsequent washes were performed using DPBS containing 0.1% Tween-20 (3 × 5 min). Slides were then incubated with the secondary antibody Alexa Fluor 488 goat anti-rabbit IgG (Invitrogen, A11008; 1:500) for 1 h at room temperature in the dark and washed again in DPBS containing 0.1% Tween-20 (3 × 5 min).

The nuclei were counterstained with 300 nM DAPI for 5 min at room temperature in the dark. The slides were washed twice with DPBS containing 0.1% Tween-20 (5 min each) and once with DPBS. Sections were mounted using antifade mounting medium and air-dried in the dark for at least 30 min, and stored at 4 °C until imaging using Evos M5000 (Invitrogen, Thermofisher).

#### Quantitative Analysis of IF

Images were analyzed using HALO v4.2 (Indica Labs) with the HighPlex FL v5.2.2 module. The thresholds for positive signal detection in each fluorescence channel were determined using the threshold-assist function and applied uniformly across all images within each marker panel. Square regions of interest (ROIs; 52 µm × 52 µm) were defined using annotation tools. The epithelial interface was manually delineated in each image. The spatial distribution of immune cells within a 200 µm peri-epithelial zone was quantified by measuring cell counts, cell density, and average distance from the epithelial interface. When multiple images or regions of interest were available from one biospecimen, ROI-level measurements were averaged to generate one value per biospecimen before statistical comparison.

#### Evaluation of spatial distribution of dendritic cells

The spatial gradient of dendritic cell density relative to the epithelial interface was quantified for NP-OPL and P-OPL biospecimens (*n* = 9 per group). For each specimen, the tissue surrounding the epithelial interface was divided into four consecutive distance bands: 0–50 μm, 50–100 μm, 100–150 μm, and 150–200 μm. The DC density within each band was calculated as the number of detected DCs divided by the corresponding band area and was expressed as cells/mm². A specimen-level density–distance slope was estimated by ordinary least-squares regression of the DC density against the midpoint of each distance band (25, 75, 125, and 175 μm). Slopes were scaled to represent the change in DC density per 50-μm increase in distance from the epithelial interface. Positive slopes indicate increasing DC density farther from the interface, whereas negative slopes indicate decreasing DC density with distance. The significance in difference in mean slopes between P-OPL and NP-OPL was evaluated using a two-sided exact permutation test by enumerating all possible reallocations of the 18 biospecimens into two groups of nine each. The exact permutation test was implemented using the Python itertools module and bootstrap confidence intervals were calculated using stratified resampling.

### Functional Analysis of S100A9

#### Cell Culture

CAL27 and FaDu cell lines were cultured in Dulbecco’s modified Eagle’s medium (DMEM) (Gibco, Cat: 11965-092) supplemented with 10% fetal bovine serum (FBS) (Corning, Cat: 35-087-CV) and 1% penicillin–streptomycin (Corning, Cat: 30-002-Cl). Immortalized human gingival keratinocytes (HGKs) (Applied Biological Materials (abm), T0717) were cultured in Prigrow X Series Medium for T0717 (abm, TM0717). Cells were maintained at 37 °C in a humidified incubator containing 5% CO₂.

#### Wound Closure (Scratch) Assay

Cells were seeded in 6-well plates and grown until they reached full confluence. A straight scratch was created across the cell monolayer using a sterile 200-µL pipette tip. Detached cells were gently removed by washing with PBS. Fresh media containing different doses of recombinant S100A9 (Cat. No. AB95909) (0, 1, and 5 µg/mL) was added. After plating, the cells were pre-treated with the TLR4 inhibitor Resatorvid (TAK-242) (100 nM, SelleckChem, Cat. No. S7455), RAGE inhibitor FPS-ZM1 (100 nM; SelleckChem, Cat. No. S8185), or an equivalent volume of DMSO, as indicated. Images of the wound area were captured at 0, 24, and 48 h using a microscope. Wound width was measured using ImageJ software. Wound closure was calculated as [(*WW*_0_ − *WW_t_*)/*WW*_0_] × 100, where *WW*_0_ is the initial wound width, and *WW_t_* is the wound width at the indicated time point.

#### Cell Proliferation Assay

Cell proliferation was assessed using the CellTrace CFSE Cell Proliferation Kit (Invitrogen Cat: C34554) according to the manufacturer’s protocol. Briefly, cells were harvested and incubated with the CFSE cell proliferation dye for 30 min at room temperature to allow intracellular labeling. Following staining, the cells were washed thoroughly with FBS to remove excess unbound dye. The labeled cells were then seeded onto culture plates and allowed to attach. After plating, the cells were pre-treated with the TLR4 inhibitor Resatorvid (TAK-242) (100 nM, SelleckChem, Cat. No. S7455), RAGE inhibitor FPS-ZM1 (100 nM; SelleckChem, Cat. No. S8185) or an equivalent volume of DMSO. Subsequently, the cells were treated with recombinant S100A8 (Cat. No. AB95343), and S100A9 (Cat. No. AB95909) recombinant protein at concentrations of 1 µg/mL and 5 µg/mL. Cells were then incubated for 72 h (cancer cells) or 96 h (human gingival keratinocytes) at 37 °C in 5% CO₂.

#### Flow Cytometry Analysis

After incubation, the cells were harvested and analyzed by flow cytometry to assess CFSE dye dilution as an indicator of cell proliferation. The CFSE fluorescence intensity decreased with each cell division, allowing quantification of proliferative activity. Data were acquired using a flow cytometer (BD FACSymphony A3 Cell Analyzer, BD Biosciences) and analyzed using the FlowJo software (FlowJo v10.1, BD Biosciences). Histograms of CFSE fluorescence intensity were generated to evaluate cell division in the treated and control cells. The primary CFSE endpoint was the division index at 72 h (cancer cells) or 96 h (human gingival keratinocytes). The technical replicate wells were averaged to produce one value for each treatment condition within each independent experiment. Three independent experiments were performed.

#### BrdU Assay

Cells were seeded onto sterile coverslips placed in 6-well plates and allowed to attach overnight. Cells were pre-treated with Resatorvid (TAK-242) TLR4 inhibitor (100 nM, Cat. No. S7455), and the FPS-ZM1 RAGE inhibitor (100 nM, Cat. No. S8185), or an equivalent DMSO vehicle for 1 h. After one hour, cells were treated with recombinant S100A8 protein (Abcam, Cat. No. AB95343) and S100A9 (Abcam, Cat. No. AB95909) at concentrations of 1 or 5 µg/mL. The cells were then incubated for 24 h at 37 °C and 5% CO2. After 24 h, cells were incubated with BrdU for 4 h. After labeling, the cells were fixed with paraformaldehyde and permeabilized with 0.3% Triton X-100 (Sigma, Cat: 024K0025). DNA was denatured using 2N HCl (Sigma, Cat: 1.09063) to expose the incorporated BrdU. Cells were blocked with 1% BSA for 1 h, followed by incubation with an anti-BrdU primary antibody (Abcam Cat: Ab6326) overnight at 4 °C. The cells were washed to remove the primary antibody and treated with Alexa Fluor 488 goat anti-rat IgG (H+L) (Invitrogen, Cat: A11006) secondary antibody for 1 h at room temperature. Cell nuclei were counterstained with DAPI (4′,6-diamidino-2-phenylindole) to visualize the total cell numbers. Fluorescent images were acquired using a fluorescence microscope (EVOS M5000; Invitrogen) under identical exposure conditions across the experimental groups. BrdU incorporation was quantified as the percentage of BrdU-positive nuclei. For each treatment condition, at least three technical wells were used for each independent experiment. The technical measurements were averaged to produce one value per treatment condition for each independent experiment. Three independent experiments were performed.

### Data Presentation and Statistical analysis

All statistical analyses were conducted using R and Python. Unless otherwise specified, all the tests were two-sided. For differential expression and pathway-enrichment analyses (DESeq2, GSEA, GO biological process enrichment), significance was defined as adjusted *p* < 0.2 after Benjamini–Hochberg false discovery rate (FDR) correction. To avoid pseudoreplication, all analyses used biospecimens as the independent biological unit. For analyses based on cell-level observations, cells were treated as nested within biospecimens, and inference was made using biospecimen-level summaries or biospecimen-clustered models, as specified.

Continuous variables were compared using the Wilcoxon rank-sum (Mann–Whitney U) test for two groups and Kruskal–Wallis test for three or more groups, followed by pairwise Wilcoxon rank-sum tests when appropriate, unless otherwise specified. Where indicated, comparisons of continuous outcomes were performed using two-sided Yuen-Welch tests with 20% trimmed means and winsorized variance estimates (allowing unequal variances between groups)—this can increase power compared to nonparametric tests, while maintaining robustness to outliers. Scratch-wound closure was analyzed using two-way repeated-measures ANOVA, with treatment and time as fixed factors and independent experiments as the repeated block, followed by Tukey’s multiple-comparison test. BrdU incorporation and CFSE-derived proliferation outcomes were measured at single endpoints and analyzed separately using one-way repeated-measures ANOVA across treatment conditions, with independent experiments as the matched block, followed by Tukey’s multiple-comparison test. Three independent experiments were analyzed, and the technical replicates were averaged for each independent experiment. Functional assay data are presented as mean ± SD.

Macrophage-subcluster abundance was analyzed using beta-binomial regression with a logit link. For each biospecimen, the response was defined as the number of macrophages assigned to the indicated sub-cluster relative to the total number of macrophages in the biospecimen. This count-based approach accounts for differences in the number of macrophages analyzed among biospecimens and allows for extrabinomial biological variability. The disease group was modeled as a categorical variable, with NP-OPL as the reference group, for pre-specified pairwise comparisons. Two-sided likelihood-ratio tests were used to calculate nominal *p* values. To evaluate ordered changes across disease states, NP-OPL, P-OPL, and OSCC were coded as 0, 1, and 2, respectively, and were included as ordinal predictors. The corresponding OR represents the change in the odds of sub-cluster membership per incremental disease stage. Because these sub-cluster analyses were exploratory, pairwise p-values were reported without adjustment for multiplicity.

Differential gene expression was tested on pseudobulk count matrices using generalized linear models with negative binomial distribution using DESeq2, with significance thresholds of FDR q<0.10 and |log2FC|≥0.25. Spatial proximity was assessed using centroid-based Euclidean distances. All pairwise epithelial–immune centroid distances were calculated within each biospecimen for the selected immune cell subsets (T cells and dendritic cells). Local epithelial neighborhoods were further defined using the k-nearest-neighbor (kNN) approach (k = 5), delineating immune neighbors within 100 µm. The neighborhood composition is summarized as the proportion of each immune cell type. All distance and neighborhood measurements were first summarized within biospecimen, and inferential analyses used one summary value per biospecimen. Pathway enrichment was performed using clusterProfiler and fgsea against the GO Biological Processes and MSigDB Hallmark sets.

To test whether S100A9-high macrophages were enriched in P-OPL compared to NP-OPL, a binomial generalized estimating equation (GEE) model was fitted with S100A9-high status as the outcome and lesion group as the predictor, using NP-OPL as the reference group. To define S100A9-high macrophages, cells were ranked according to S100A9 expression across the full macrophage dataset, and cells with S100A9 values greater than the cohort-wide median value were classified as S100A9-high. The biospecimen ID was specified as the clustering variable with an exchangeable working correlation structure to account for the non-independence of cells derived from the same biospecimen. Wald test was used for statistical significance.

For predictive model build-up and validation of the markers, malignant progression was modeled as a binary patient-level outcome. Progressive lesions were defined as premalignant lesions that developed oral squamous cell carcinoma within 5 years of the index biopsy, whereas non-progressive lesions showed no malignant transformation during at least 5 years of follow-up. The modeling cohort included 23 patients, comprising 11 progressive and 12 non-progressive lesions. The expanded clinical candidate set comprised age, sex, histologic grade dichotomized as mild versus non-mild, ordinal tobacco exposure coded as 1=never, 2=former, and 3=current, a tobacco-missingness indicator, and a pre-specified high-risk versus low-risk anatomical site variable. Alcohol exposure was excluded a priori because 8 of the 23 values were missing. The expanded clinical– biomarker candidate set comprised six clinical variables and all eight available biomarker readouts: MX1 cytoplasmic intensity, MX1 cell intensity, S100A9 cytoplasmic intensity, S100A9 cell intensity, mean count of TIGIT^+^ T cells in 200 μm of epithelial dysplastic interface, median count of TIGIT^+^ T cells in 200 μm of epithelial dysplastic interface, TIGIT intensity mean, and TIGIT intensity median. Candidate predictors were evaluated using elastic net–penalized logistic regression. The mixing parameter (*l1_ratio*) was evaluated over 0.1, 0.5, 0.9, and 1.0, and the inverse regularization-strength parameter (C) over 0.01, 0.03, 0.1, 0.3, and 1.0. Hyperparameters were selected by stratified fivefold cross-validation optimizing log-loss using the one-standard-error rule, with ties favoring a higher *l1_ratio* to encourage parsimony. The full-data clinical model selected *l1_ratio*=1.0, and C=0.3, whereas the clinical–biomarker model selected *l1_ratio*=0.1, and C=0.1. Selected predictors were refitted using *L*_2_-penalized ridge logistic regression with default value of C=1.

Within each resampling split, missing continuous values were imputed using the predictor-specific median calculated across all patients in the training partition without stratification by progression status. Continuous predictors were standardized using training set parameters. The complete preprocessing, hyperparameter-tuning, variable selection, and refitting pipeline were repeated in 1,000 stratified bootstrap resamples and in 50 repetitions of nested five-fold cross-validation; both *l1_ratio* and C were therefore re-tuned within each resample or training fold. Because *l1_ratio*=0.1 permits correlated biomarkers to share coefficients, the recurrent clinical plus biomarker core was defined using bootstrap selection frequencies. Histologic grade (mild, non-mild), S100A9 cytoplasmic intensity, and mean count of TIGIT^+^ T cells in 200 μm of epithelial dysplastic interface were selected in 76.6%, 70.8%, and 67.5% of the bootstrap resamples, respectively. These three predictors were then evaluated as a fixed panel using ridge logistic regression with default value C=1.0.

A separate, hypothesis-driven clinical–spatial model was evaluated to assess added value of spatial features using a prespecified reduced candidate set comprising age, sex, histologic grade, ordinal tobacco exposure, tobacco-missingness indicator, dendritic-cell average distance from the epithelial interface, log1p-transformed dendritic cell density, CD3-positive T-cell average distance from the epithelial interface, and TIGIT-positive T cell average distance from the epithelial interface. Dendritic-cell density was log1p-transformed because of the marked right skew, whereas distance measurements were retained on the original micrometer scale. A TIGIT-positive T-cell distance of zero, accompanied by zero TIGIT-positive T-cell density, was treated as the absence of the measured population and coded as missing. Variable selection was performed using *L_1_*-penalized logistic regression tuned by stratified five-fold cross-validation with the one-standard-error rule. The retained predictors were refitted using ridge-stabilized logistic regression with default value of C=1.0. Full-pipeline bootstrap and repeated nested cross-validation were performed, with preprocessing, variable selection, and model fitting repeated within each training sample.

Model discrimination was summarized using the area under the receiver operating characteristic curve (AUC), and the overall prediction error was summarized using the Brier score. We report apparent and bootstrap optimism-corrected values.

The analyses were performed using R (v4.4.1) and Python (v3.13.5). Major R packages included Seurat (v5.4.0) and SingleR (v2.6.0) with Celldex (v1.14.0) for cell type annotation; DESeq2 (v1.44.0) for differential expression analysis; clusterProfiler (v4.12.6), fgsea (v1.30.0), and enrichplot (v1.24.4) with msigdbr (v26.1.0) for gene set enrichment and pathway analysis; and nichenetr (v2.2.0) for ligand-receptor interaction analysis. The dendritic-cell spatial gradient analysis was performed in Python (v3.12.13) using pandas (v2.2.3), NumPy (v2.3.5), and SciPy (v1.17.0) for numerical calculations and linear regression, and Matplotlib (v3.10.8) for visualization. The clinical–spatial model was fitted in Python (v3.13.5) using NumPy (v2.3.5), SciPy (v1.17.0), scikit-learn (v1.8.0), and Joblib (v1.5.3), with random seed 20260726. Xenium data were inspected using Xenium Explorer and processed using Xenium Onboard Analysis. Immunofluorescence images were analyzed using HALO v4.2 with HighPlex FL v5.2.2.

## RESULTS

### Spatial transcriptomic profiling of NP-OPL, P-OPL, and OSCC samples

To resolve cell-type heterogeneity and disease-stage transcriptional programs, we applied single-cell-resolution Xenium spatial transcriptomics to 20 HPV-negative biospecimens from 16 patients: eight nonprogressive oral premalignant lesions (NP-OPLs), eight progressive OPLs (P-OPLs) collected before malignant transformation, and four subsequently developed OSCC specimens longitudinally matched to four of the P-OPLs (**Fig. 1A; Fig. S1A; Table S1**). Thus, OSCC specimens did not constitute an unrelated cross-sectional cancer cohort: each was a follow-up specimen from a patient whose antecedent P-OPL was included in the analysis. This matched design provided a longitudinal context for the transition from premalignancy to invasive disease and reduced between-patient confounding in comparisons between P-OPLs and OSCC, although the number of matched pairs was limited. After quality control, all 20 biospecimens were retained, yielding 523,753 cells and 41,326,882 decoded transcripts for downstream analysis. Xenium output was reviewed in Xenium Explorer 3 for segmentation quality and exported as spatial experimental files and 10x-format matrices. The matrices were analyzed using Seurat v5, and the corresponding tissue images and spatial coordinates were used for in situ analysis.

**Fig. 1.**
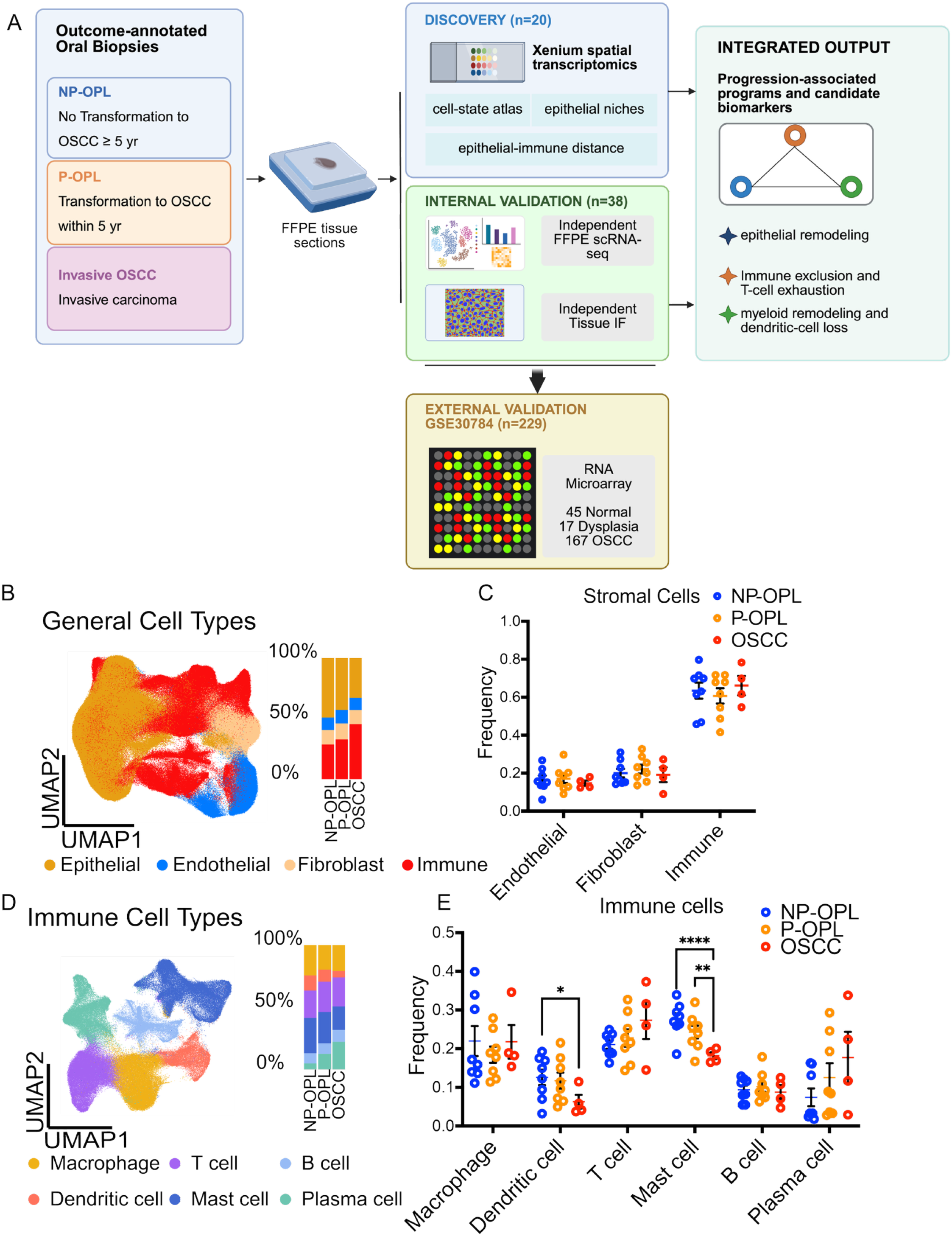
Characterization of FFPE-preserved NP-OPL, P-OPL and OSCC biospecimens by Xenium spatial transcriptomics. Analysis of 20 biospecimens identified heterogeneity in cell composition across non-progressive oral premalignant lesions (NP-OPL) (n=8), progressive oral premalignant lesions (P-OPL) (n=8) and oral squamous cell carcinoma (OSCC) (n=4). **A** Schematic illustration of the study workflow. **B** UMAP visualization of major cell types consolidated from Seurat clusters, including epithelial cells, fibroblasts, endothelial cells and immune cells, annotated using canonical marker expression. **C** Biospecimen-level frequencies of major cell types across disease groups. **D** UMAP visualization of immune cell types, including T cells, macrophages, plasma cells/immunoglobulin-expressing cells, mast cells, B cells and dendritic cells. **E** Relative frequencies of macrophages, dendritic cells, T cells, mast cells, B cells, and plasma cells within the total immune-cell population across biospecimens. Each point represents one biospecimen. Pairwise differences between disease groups were assessed using two-sided Yuen’s tests, with standard errors estimated from winsorized variances. *, **, *** and **** indicate *p* < 0.05, *p* < 0.01, *p* < 0.001 and *p* < 0.0001, respectively. Data are presented as mean ± SEM.

### Global cell type composition across disease states

The number of principal components to use for PCA dimensionality reduction was selected using an elbow plot of the first 20 PCs; 15 PCs were retained (**Fig. S1B–C)**. Cells were clustered using the Louvain algorithm and embedded in UMAP using these PCs. We identified 15 clusters, which were consolidated into four major lineages by canonical markers: epithelial cells (*EGFR*, *EPCAM*, *ERBB2/3*, *CEACAM*s), fibroblasts (*FN1*), endothelial cells (*PLVAP*, *SPARCL1*), and immune cells, with the latter comprising T cells (*IL2RB*, *TRAT1*), macrophages (*MPEG1*, *CD163*), plasma cells (immunoglobulin), mast cells (*CPA3*, *P2RX1*), B cells (*MS4A1*, *PAX5*), and dendritic cells (*CD1A*, *CD1C*) (**Fig. S2**). Marker expression for each consolidated lineage was visualized to confirm the identity assignments (**Fig. S3; Tables S4 and S5**). Cell type and subcluster frequencies are expressed as the proportion of cells within their parent compartment. We compared the frequency of major cell types among NP-OPL, P-OPL, and OSCC biospecimens and detected no significant differences across disease states (**Fig. 1B, C; Fig. S4**).

### Immune cell dynamics in progression to OSCC

To further resolve the immune composition, we analyzed the individual immune cell types **(Fig. 1D; Fig. S4)**. Analysis of the major immune cell populations demonstrated changes in the immune composition associated with the progression to OSCC. Dendritic cell frequency was lower in OSCC than in NP-OPL (p<0.05). Mast-cell frequency was also lower in OSCC than in both NP-OPL (p<0.0001) and P-OPL (p<0.01). As a complementary biospecimen-level analysis, the mast-cell frequency decreased monotonically across NP-OPL, P-OPL, and OSCC (28.0%, 24.3%, and 18.2%, respectively; two-sided Cuzick test for trend, *p* < 0.01) **(Fig. 1E)**. This trend remained significant after false discovery rate correction across the six immune cell populations (q<0.05). No significant ordered trend was detected in other immune cell populations. These findings support the progressive loss of mast cells during malignant transformation, together with reduced dendritic cell representation in OSCC. To capture more granular heterogeneity and identify potential progression-associated biomarkers, we performed sub-clustering of epithelial cells, macrophages, T cells, and dendritic cells. This analysis revealed distinct subpopulations with differential abundances across disease stages, suggesting that immune remodeling, rather than global epithelial or stromal shifts, represents a key feature of malignant progression.

### Sub-clustering of epithelial cells reveals progression-associated states

Eight epithelial subclusters were identified **(Fig. 2A)**, and their marker profiles and relative abundances were compared across the NP-OPL, P-OPL, and OSCC biospecimens **(Fig. 2B, C, Fig. S5; Table S6)**. Epi.sub.1 frequency was increased in both P-OPL (p<0.05) and OSCC (p<0.01) compared to NP-OPL **(Fig. 2B)**. Marker analysis identified characteristic genes for each epithelial sub-cluster, with the selected marker sets visualized in **Fig. 2C**. This subcluster represents a progression-associated epithelial state marked by *MX1, NOTCH3, PTGS1* (*COX-1*), and *NELL2*. This signature is consistent with type-I interferon/stress signaling (*MX1*), *NOTCH3*-linked epithelial remodeling, and prostaglandin biosynthesis (*PTGS1*).

**Fig. 2.**
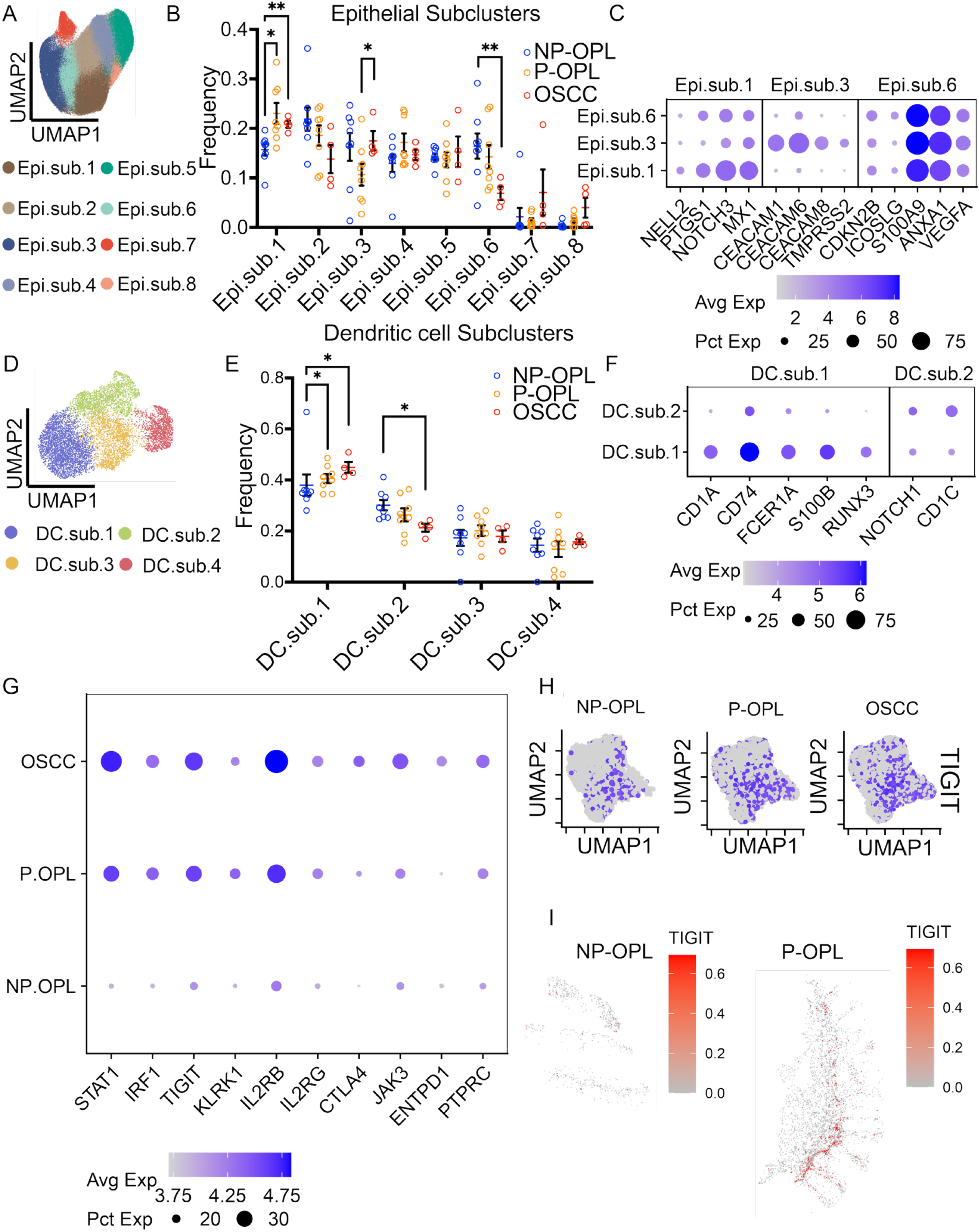
Subcluster characterization of epithelial cells, dendritic cells and T cells across disease states. **A** UMAP visualization of eight epithelial subclusters in Xenium data set (n=20 biospecimens). **B** Biospecimen-level frequencies of epithelial subclusters across disease groups. **C** Marker expression of epithelial subclusters exhibiting significant disease-associated frequency changes, including Epi.sub.1. **D** UMAP visualization of four dendritic cell subclusters. **E** Biospecimen-level frequencies of dendritic-cell subclusters across disease groups. **F** Marker expression of dendritic-cell subclusters exhibiting disease-associated frequency changes, including DC.sub.1 and DC.sub.2. **G** Differentially expressed T-cell markers in P-OPL and OSCC compared with NP-OPL. **H** Feature plots showing increased *TIGIT* expression across progressive lesions and OSCC. **I** Representative Xenium image showing *TIGIT* enrichment in P-OPL. Prespecified pairwise comparisons were performed using two-sided Yuen tests. * and ** indicate *p* < 0.05 and *p* < 0.01, respectively. Data are presented as mean ± SEM. Individual biospecimen-level values are shown.

The frequency of Epi.sub.3, marked by enriched expression of *TMPRSS2, CEACAM8, CEACAM6,* and *CEACAM1*, was increased in OSCC compared to P-OPL (p<0.05) **(Fig. 2B, C; Table S6)**. Enrichment of *CEACAM* family genes and *TMPRSS2* is consistent with a barrier/secretory epithelial program that increases during progression. CEACAM-family molecules have been implicated in mucosal epithelial and neutrophil biology, and CEACAM6 promotes OSCC invasion and metastasis through complex N-glycosylation-dependent EGFR signaling (*22, 23*). TMPRSS2 has been detected in oral epithelial tissues and is embedded in the IL-13-associated mucus secretory programs in the airway epithelium (*24, 25*). Together, these features suggest that Epi.sub.3 represents an epithelial state that is crucial for malignant transformation.

Epi.sub.6 frequency reduced in OSCC compared with NP-OPL (p<0.01) and it showed enriched expression of *ICOSLG* and *CDKN2B* compared to other subclusters but lacked a single uniquely expressed marker with definitive lineage or function **(Fig. 2B, C; Table S6)**, consistent with the loss of an epithelial state with potential local immunomodulatory or costimulatory functions during progression.

Complementary ordered-trend analysis identified an association between Epi.sub.1 frequency and disease stage (16.2%, 22.5%, and 20.9% in NP-OPL, P-OPL, and OSCC, respectively; Cuzick p<0.01; q<0.01). Because Epi.sub.1 frequency increased from NP-OPL to P-OPL and remained similarly elevated in OSCC, this finding represents an ordered association rather than a strictly stepwise increase.

GO enrichment analysis of epithelial sub-cluster markers further supported distinct functional programs among progression-associated epithelial states. Epi.sub.1 markers were enriched for cytokine-mediated signaling and regulation of cell population proliferation, Epi.sub.3 markers were enriched for inflammatory and cytokine-production programs, and Epi.sub.6 markers were enriched for epithelial proliferation, cell survival, and cytokine/T-cell regulatory terms **(Fig. S6; Tables S7-S9).**

### Dendritic cell subclusters and functional constraints in progression

We resolved four dendritic cell (DC) sub-clusters and compared their marker profiles and frequencies across NP-OPL, P-OPL, and OSCC samples **(Fig. 2D-F; Fig. S7; Table S10)**. The frequency of DC.sub.1, marked by *CD1A, CD74, FCER1A, S100B* and *RUNX3*, was increased in P-OPL (p<0.05) and OSCC (p<0.05) despite the overall contraction of the DC compartment **(Fig. 1E and Fig. 2E, F)**. This profile is consistent with a tissue-resident, antigen-processing, conventional DC type 2 (cDC2)/Langerhans-like state. GO enrichment analysis of DC.sub.1 markers showed enrichment for cytokine-mediated signaling, cytokine production, and microbial-response pathways **(Fig. S8; Table S11)**.

In contrast, DC.sub.2, marked by *NOTCH1* and *CD1C*, its frequency decreased in OSCC (p<0.05) **(Fig. 2E, F)**. This pattern is consistent with the loss of Notch1-associated cDC2 program linked to CD4+ T-cell priming (Table S12). Together, these findings suggest that DC remodeling during progression involves selective retention of tissue-resident DC-like states, alongside the loss of DC programs associated with T-cell priming and epithelial immune surveillance. DC.sub.1 progressively increased across NP-OPL, P-OPL, and OSCC (34.9%, 40.3%, and 44.9%; Cuzick p<0.05, q<0.05). Conversely, DC.sub.2 progressively decreased (29.3%, 26.5%, and 21.3%; Cuzick p<0.05, q<0.05). No ordered trends were detected for DC.sub.3 or DC.sub.4.

### TIGIT-high, exhaustion-associated T-cell states increase in progressive lesions and OSCC

We identified elevated expression of markers of T cell in P-OPL and OSCC relative to NP-OPL **(Fig. 2G; Fig. S9; Tables S13 and S14)**. Among these markers, *TIGIT* showed a progressive increase across disease states from NP-OPL to P-OPL to OSCC **(Fig. 2H)**, suggesting the enrichment of inhibitory, exhaustion-associated T-cell programs during malignant progression. This pattern was further supported by the spatial expression analysis **(Fig. 2I)**. GO enrichment analysis identified distinct T-cell pathway enrichment in P-OPL and OSCC, including terms related to T-cell activation and cytokine-mediated signaling, consistent with dysregulated T-cell activation during progression **(Fig. S10; Tables S15 and S16).**

### Macrophage heterogeneity and inflammatory remodeling in OSCC

Macrophages were sub-clustered from the immune compartment, yielding four sub-clusters **(Fig. 3A, B, Fig. S11A, and Table S17)**. Mac.sub.3 was characterized by *APOE*, *MMP12*, *S100A9*, *ISG15*, and *SDC1* **(Fig. 3C)**. In the ordered beta-binomial model, Mac.sub.3 increased across NP-OPL, P-OPL, and OSCC, with a 1.78-fold increase in the odds of Mac.sub.3 membership per incremental disease stage (two-sided likelihood-ratio, p<0.01). In pairwise beta-binomial analyses, the odds of Mac.sub.3 membership were higher in OSCC than in NP-OPL (p<0.01) and P-OPL (p<0.05). The difference between the P-OPL and NP-OPL groups was not significant (p>0.05) **(Fig. 3B, C)**. This population was consistent with an inflammatory remodeling and interferon-responsive macrophage phenotype. *S100A9* and *ISG15* are associated with innate activation and type-I interferon stress (*26–29*), *MMP12* is linked to matrix remodeling and tissue invasion (*30*), and *APOE*/*SDC1* has been implicated in lipid-ECM programs associated with immunosuppressive myeloid reprogramming in tumors (*31–34*). GO enrichment analysis of Mac.sub.3 markers further supported the cytokine-mediated signaling pathway and positive regulation of cell population proliferation in this macrophage subcluster **(Fig. S11B; Table S18)**.

**Fig. 3.**
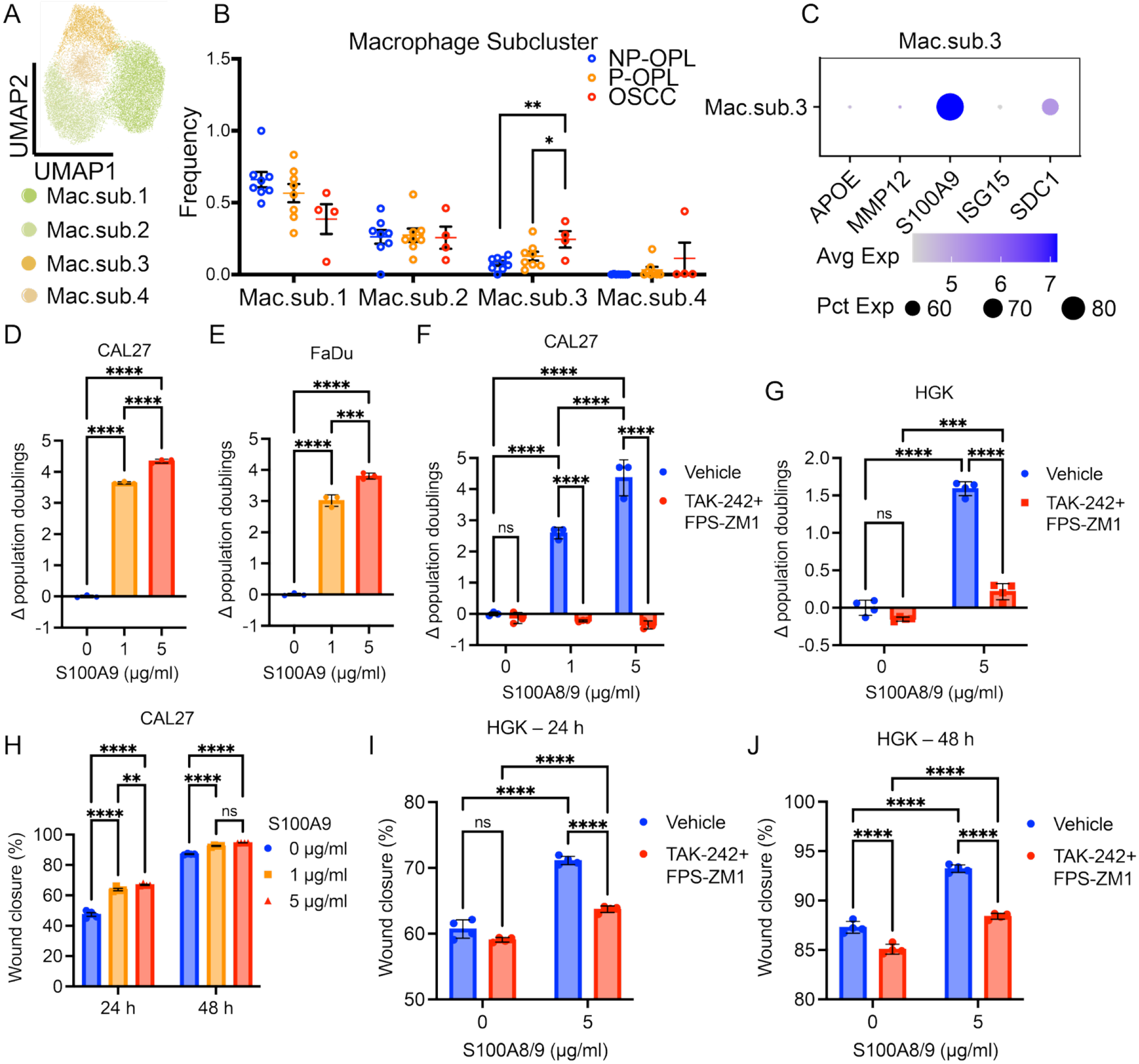
A S100A9-high macrophage subcluster is enriched in oral cancer; recombinant S100A8/9 promoted cell proliferation and migration via TLR4 and RAGE in CAL27, FaDu, and human gingival keratinocytes (HGKs) in vitro. **A** UMAP visualization of macrophage (Mac) subclusters in Xenium dataset (n=20 biospecimens). **B** Macrophage-subcluster abundance was calculated as the number of macrophages assigned to the indicated subcluster relative to the total number of macrophages. Differences among groups were evaluated using beta-binomial regression with a logit link. Pairwise comparisons are annotated with asterisks corresponding to nominal *p*-values. Data are presented as mean ± SEM. **C** Dot plot showing marker expression in Mac.sub.3. **D-E** S100A9 induced cells division in CAL27 (**D**) and in FaDu (**E**) cells. One-way ANOVA with Tukey’s correction for multiple comparisons were used for statistical analysis. **F** The TLR4 inhibitor TAK-242 and the RAGE inhibitor FPS-ZM1 reduced S100A8/9-induced cell division in CAL27 cells in vitro. Two-way ANOVA with Tukey’s correction for multiple comparisons were used for statistical analysis. **G** TAK-242 and FPS-ZM1 inhibited S100A8/9–induced proliferation in HGK in vitro. Δ population doublings are values calculated as division index of each group minus the mean in vehicle + 0 µg/ml group. Two-way ANOVA with Bonferroni’s correction for multiple comparisons was used for statical analysis. **H** S100A9 promoted wound closure in CAL27 cells. **I-J** TAK-242 and FPS-ZM1 inhibited S100A8/9-induced wound closure in HGKs at 24 h (**I**) and at 48 h (**J**). Wound closure was calculated as the reduction in wound width from baseline, expressed as a percentage of the initial width. Two-way ANOVA with Tukey’s correction for multiple comparisons was used for statistical analysis. (**D–J)** data are presented in mean ± SD. * *p* < 0.05, \*\**p* < 0.01, \*\*\**p* < 0.001, \*\*\*\**p* < 0.0001; ns, not significant.

### S100A9-associated wound-closure and proliferative responses are attenuated by pharmacologic inhibition of TLR4 or RAGE

To determine whether S100A9 influences wound closure in oral cancer cells, we performed CellTrace CFSE cell proliferation assay and wound closure assays in OSCC cells (CAL27 and FaDu) and human gingival keratinocytes (HGKs). To validate the functional effects associated with S100A9, we used a combination of recombinant S100A8 and S100A9, which are predominantly associated with the heterodimeric complex calprotectin in myeloid cells, and their extracellular signaling activity is strongly influenced by complex formation and the oligomeric state (*35, 36*). The combination of recombinant S100A8 and S100A9 is a well-characterized, biologically active inflammatory ligand reported to engage pattern-recognition pathways associated with TLR4 and RAGE (*35, 36*). Thus, treatment with a combination of recombinant S100A8 and S100A9 provides a biologically relevant approach to determine whether activation of the S100A9-containing calprotectin axis could reproduce the phenotype identified in our discovery analysis. Treatment with S100A9 or recombinant S100A8/9 proteins promoted cell divisions in CAL27, FaDu, and HGKs **(Fig. 3D–G; Fig. S12-14)**. S100A9 treatment alone was sufficient to accelerate wound closure compared to untreated controls in CAL27 cells while S100A8/9 treatment did so in HGKs, both with greater closure observed at 24 and 48 h after treatment **(Fig. 3H–J; Fig. S12, 14**).

To test whether this effect was mediated through canonical S100A9-associated receptors, cells were treated with TAK-242, a TLR4 inhibitor, and FPS-ZM1, a RAGE inhibitor. Pharmacological inhibition of TLR4 and RAGE reduced S100A8/9-induced cell divisions in CAL27 cells and HGKs, suggesting that these receptors may contribute to the proliferative response **(Fig. 3F, G; Fig. S12, 14)**. Inhibition of TLR4 and RAGE also attenuated S100A8/9-induced wound closure in CAL27 cells and HGKs (**Fig. 3H–J**), indicating that S100A8/9 promotes epithelial cell migration and proliferation simultaneously, at least in part, through TLR4- and RAGE-associated signaling, consistent with previous reports linking S100A8/9 inflammatory activity to these receptor pathways (*36*). Furthermore, we tested whether S100A8/9 could play a role in facilitating DNA synthesis. In CAL27 cells, S100A8/9 treatment increased BrdU incorporation, indicating S100A8/9 may contribute to enhanced DNA synthesis and thereafter cell division **(Fig. S12D)**. These findings indicate that S100A9 enhances the capacity of cell proliferation and migration in both cancer and normal epithelial cells, and, collectively, support a pro-proliferative role of S100A9 signaling in oral epithelial and OSCC cells.

### Spatial organization of epithelial-immune cell interactions

Spatial proximity analysis further demonstrated an increased separation between basal epithelial cells and selected immune populations during progression. Compared with NP-OPL, T cells and dendritic cells in P-OPL were positioned approximately 22 and 11 μm farther from the epithelial basal layer, respectively. In OSCC, T cells and dendritic cells were positioned approximately 16 μm away from the epithelial basal layer relative to the NP-OPL **(Fig. 4A).**

**Fig. 4.**
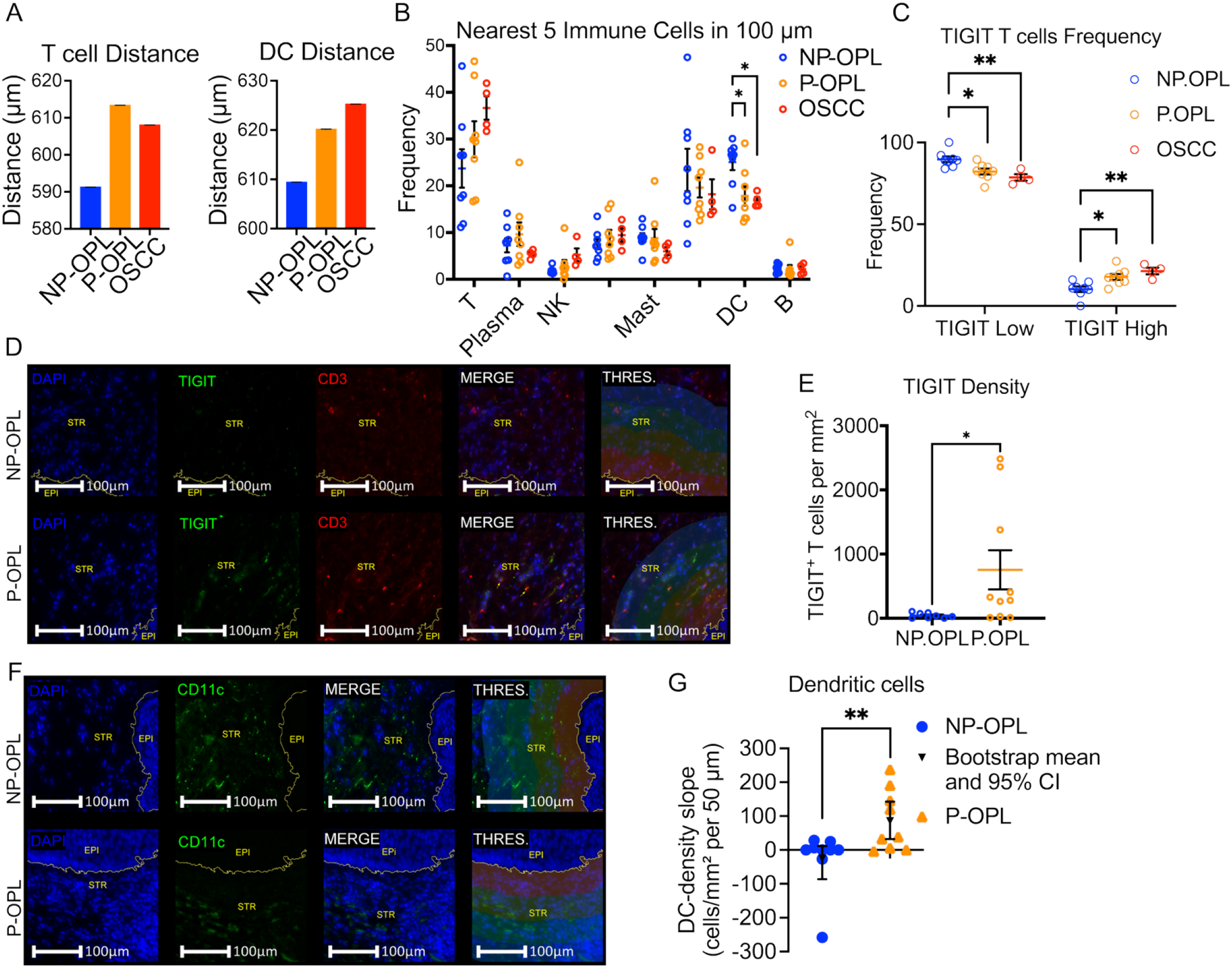
Spatial organization of epithelial-immune interactions reveals immune remodeling in P-OPL and OSCC. **A** Distances from the epithelial basal layer to T cells and dendritic cells across NP-OPL, P-OPL, and OSCC. **B** Frequencies of immune-cell types among the five nearest immune-cell neighbors located within 100 μm of each basal epithelial cell. **C** Frequencies of TIGIT-low and TIGIT-high T cells among the two nearest T cell neighbors located within 100 μm of each basal epithelial cell. For panels B–C, each point represents one independent Xenium biospecimens (n = 20). **D** Representative immunofluorescence images and corresponding HALO analyses of CD3⁺TIGIT⁺ T cells within a 200-μm epithelial-niche region (THRES.; four 50-µm intervals annotated) extending from the epithelial interface into the underlying stroma. Scale bars, 100 μm. **E** Quantification of CD3⁺TIGIT⁺ T-cell density within the epithelial-niche region in NP-OPL and P-OPL biospecimens (n = 8 NP-OPL and n = 10 P-OPL). **F** Representative immunofluorescence images (with slope near mean) and corresponding HALO analyses of CD11c⁺ dendritic cells within a 200-μm epithelial-niche region (THRES.; four 50-µm intervals annotated) extending from the epithelial interface into the underlying stroma. Scale bars, 100 μm. **G** Specimen-level gradients of DC density were calculated across four consecutive distance bands from the epithelial interface (0–50, 50–100, 100–150, and 150–200 μm). Each point represents the density– distance slope for one independent biospecimen (n = 9 NP-OPL and n = 9 P-OPL), expressed as the change in DC density per 50-μm increase in distance. Triangles and error bars indicate group means and bootstrap 95% confidence intervals, respectively; the horizontal line denotes a slope of zero. (**B–C)** overall differences among NP-OPL, P-OPL, and OSCC were assessed using Kruskal–Wallis tests, followed by two-sided Mann–Whitney U tests for pairwise comparisons between disease groups. (**E)** NP-OPL and P-OPL were compared using a two-tailed Mann–Whitney test. (**G**) The difference between NP-OPL and P-OPL was assessed using a two-sided exact permutation test. (**A-C, E**) Data are presented as mean ± SEM. \**p* < 0.05, \*\**p* < 0.01.

Next, we examined the immune cell composition within the epithelial niche in the Xenium dataset, defined as non-epithelial cells located within 100 μm of the basal epithelial layer. Among the five nearest immune cell neighbors to each basal epithelial cell, dendritic cells were reduced in OSCC relative to NP-OPL, whereas TIGIT-high (at least 1 TIGIT transcript detected) T cells were increased in OSCC and TIGIT-low (no TIGIT detected) T cells were decreased **(Fig. 4B, C)**. These findings indicate the remodeling of the immune compartment adjacent to the epithelial interface.

To validate spatial immune remodeling, we performed immunofluorescence staining for TIGIT-positive T cells in an independent cohort of NP-OPL (n=8) and P-OPL (n=10) biospecimens. CD3-positive TIGIT-positive T cells were quantified within a 200 μm region extending from the epithelial layer toward the stroma **(Fig. 4D)**. Consistent with the Xenium finding of increased TIGIT-high T cells near basal epithelial cells, immunofluorescence analysis showed higher CD3-positive TIGIT-positive T-cell density within the 200 μm epithelial niche region of P-OPL biospecimens than NP-OPL biospecimens (p<0.05) **(Fig. 4E)**. These findings support the enrichment of TIGIT-associated T cell states in the P-OPL epithelial microenvironment.

To verify the spatial immune changes involving DCs, we quantified CD11c-positive DCs by immunostaining in an independent cohort (n=9 NP-OPL; n=9 P-OPL). DC density demonstrated significantly different spatial gradients in the P-OPL and NP-OPL biospecimens **(Fig. 4F, G)**. NP-OPL biospecimens generally exhibited negative density–distance slopes, consistent with higher DC density near the epithelial interface and decreasing density at greater distances. In contrast, P-OPL biospecimens exhibited more positive slopes, indicating redistribution of DCs toward regions farther from the epithelial interface. The mean slope was 110.4 cells/mm² per 50 μm higher in P-OPL than in NP-OPL (bootstrap 95% CI=38.8–194.2; two-sided exact permutation, p<0.01) **(Fig. 4G)**. These findings support the progressive displacement of DCs away from the epithelial interface of the P-OPL.

### Single-cell validation in an independent cohort

To validate the spatial transcriptomic findings in an independent patient cohort, we performed whole-transcriptome single-cell RNA sequencing using the 10x Genomics scRNA Flex workflow on 16 FFPE oral premalignant lesion biospecimens from 16 unique patients, including 11 progressive OPLs (P-OPLs) and five non-progressive OPLs (NP-OPLs). Each patient contributed one specimen. This cohort was analyzed independently from the Xenium discovery cohort and was used to test whether progression-associated epithelial, myeloid, dendritic, and T-cell signatures identified by spatial transcriptomics were reproducible at whole-transcriptome single-cell resolution.

After quality control, 54,096 nuclei were retained for validation analysis, and one sample from the P-OPL failed to pass QC metrics. Major lineages were recovered using canonical markers, including epithelial cells (*KRT5, KRT14, S100A2, CSTA, SPRR1B*), endothelial cells (*ACKR1, RAMP2, SELE, VWF, PECAM1*), fibroblasts (*LUM, COL3A1, DCN, COL1A1, CFD*), and immune cells (*CD69, CD52, CXCR4, PTPRC, HCST*), consistent with previous oral mucosa atlases (*8*). Additional compartments included the muscle, salivary gland cells, and melanocytes **(Fig. 5A; Fig. S15A, Fig. S16; Table S19)**. Immune cells were further subtyped using SingleR with the Monaco immune reference (*15*) and verified by marker expression **(Fig. 5B; Fig. S15B; Fig. S16; Table S20).**

**Fig. 5.**
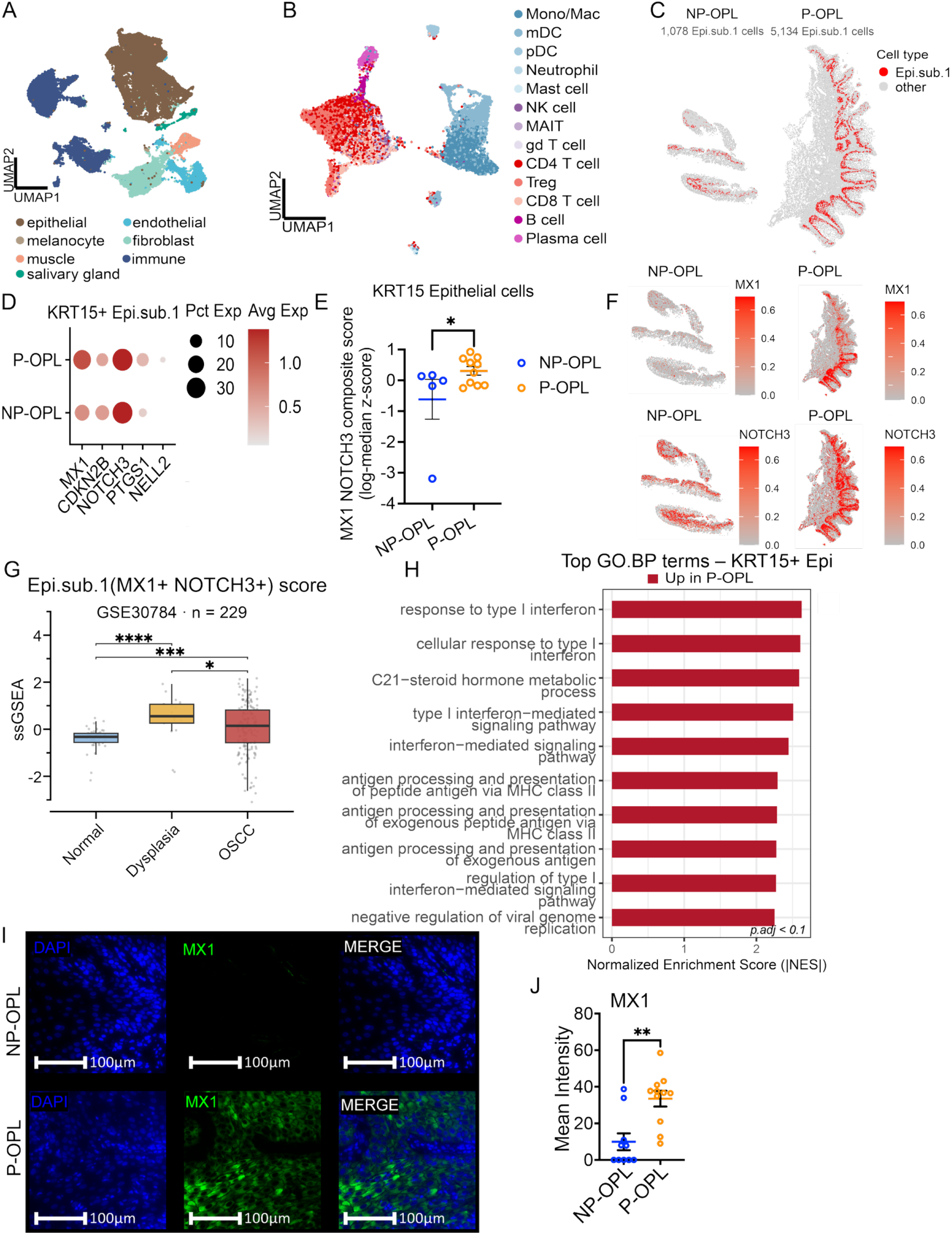
Independent whole transcriptomic scRNA-seq validation of progression-associated epithelial programs. **A** UMAP of major cell types identified in an independent scRNA-seq cohort in 15 independent biospecimens from 15 unique patients (5 NP-OPL and 10 P-OPL). **B** UMAP visualization of immune cell types annotated using SingleR with the Monaco immune reference. **C** Representative Xenium image plot showing basally enriched Epi.sub.1 population in NP-OPL and P-OPL. **D** Dot plot showing expression of KRT15+ epithelial markers. **E** Biospecimen-level composite score for *MX1* and *NOTCH3* expression in KRT15+ epithelial cells. Each point represents one independent biospecimen; statistical significance was assessed using a two-sided exact permutation test at the specimen level. Data are presented as mean ± SEM. **F** Representative Xenium feature plots showing *MX1* and *NOTCH3* expression in NP-OPL and P-OPL biospecimens. **G** Box plot of single-sample gene set enrichment analysis (ssGSEA) scores of Epi.sub.1 markers in **D** in normal, dysplasia, and OSCC samples showed score enrichment along with malignant progression (GSE30784; n = 229; 45 normal/ 17 dysplasia/ 167 OSCC). Groups were compared by two-tailed Mann–Whitney tests across all three pairwise contrasts with Benjamini–Hochberg correction. \**q* < 0.1, \*\*\**q* < 0.001, \*\*\*\**q* < 0.0001. **H** GSEA of Gene Ontology Biological Process gene sets in KRT15+ epithelial cells. **I** Representative immunofluorescence images of MX1 staining in epithelial regions. Scale bars, 100 μm. **J** Quantification of epithelial MX1 mean fluorescence intensity (n=10 for NP-OPL and n=11 for P-OPL). NP-OPL and P-OPL were compared using a two-tailed Mann–Whitney test. \**p* < 0.05, \*\**p* < 0.01, \*\*\**p* < 0.001, \*\*\*\**p* < 0.0001.

#### Epithelial validation

Spatial mapping in the Xenium dataset localized Epi.sub.1 cells to the basal epithelial layer, within regions characterized by *KRT14* and *KRT15* expression **(Fig. 5C)**. We subsetted epithelial cells in the independent scRNA-seq cohort to examine the basal *KRT14*+ and *KRT15*+ compartments, as these compartments lie at the epithelial–stromal interface and directly interact with infiltrating immune cells. In *KRT15*+ epithelial cells, progression-associated markers identified in the Xenium analysis, including *MX1* and *NOTCH3,* showed higher expression in P-OPL than in NP-OPL (p<0.05) **(Fig. 5D, E; Fig. S17A)**. Spatial feature plots in representative Xenium images further showed increased *MX1* and *NOTCH3* expression in the P-OPL epithelial regions **(Fig. 5F)**. Similar marker expression patterns were observed in *KRT14*+ epithelial cells in the validation cohort **(Fig. S18)**. Validation in a published external cohort (GSE30784; n = 229; 45 normal/ 17 dysplasia/ 167 OSCC) confirmed the elevated ssGSEA score of Epi.sub.1 markers (*MX1, CDKN2B, NOTCH3, PTGS1, NELL2*) along with disease progression from normal to dysplasia and thereafter to OSCC **(Fig. 5G)**, with specific enrichment in dysplasia, suggesting its unique role in dysplastic tissues. We performed a pseudobulk analysis on *KRT15*+ and *KRT14*+ cells in P-OPL vs. NP-OPL (n=15; 5 NP-OPL, 10 P-OPL) **(Table S21, Table S22)**. Pathway and GSEA analyses of *KRT14*+ and *KRT15*+ epithelial subsets supported the enrichment of inflammatory and stress-response programs in P-OPL, with type-I and type-II interferon response pathways enriched in both basal epithelial compartments **(Fig. 5H; Fig. S19; Tables S23-S26)**. These findings validated a progression-associated basal epithelial inflammatory program across independent spatial and single-cell datasets. Immunofluorescence staining in an independent cohort of 21 biospecimens from 21 unique patients, including 10 NP-OPL and 11 P-OPL biospecimens/patients, was used to validate the MX1 protein levels in epithelial cells. HALO-based image analysis identified an increased intensity of MX1 in epithelial cells in P-OPL versus NP-OPL **(Fig. 5I, J)**.

#### Macrophage validation

P-OPL samples contained a higher fraction of S100A9-high macrophages than NP-OPL samples, (p<0.05) **(Fig. 6A; Fig. S17B)**. Spatial feature plots from representative Xenium images revealed enrichment of *S100A9* signals in P-OPL samples **(Fig. 6B)**. Validation of the ssGSEA score of S100A9+ Mac.sub.3 markers (*APOE, MMP12, S100A9, ISG15, SDC1*) in the external cohort (GSE30784; n = 229; 45 normal/ 17 dysplasia/ 167 OSCC) showed a progression-associated elevation of the score, which monotonically increased from normal to dysplasia and to OSCC **(Fig. 6C).** We performed a pseudobulk analysis on macrophages in P-OPL vs. NP-OPL (n=15; 5 NP-OPL, 10 P-OPL) **(Table S27)**. GSEA and pathway analyses supported the upregulation of TNF-α/NF-κB, interferon-α/γ, and inflammatory response programs in the P-OPL macrophages **(Fig. 6D; Table S28)**. GO biological process analysis further identified the enrichment of T-cell apoptotic processes, antimicrobial humoral response, granulocyte migration, and toll-like receptor signaling pathways in macrophages in P-OPL **(Fig. 6E; Table S29)**. Immunofluorescence staining in an independent cohort of 22 biospecimens from 22 unique patients, including 12 NP-OPL and 10 P-OPL biospecimens/patients, was used to validate the S100A9 protein levels near the epithelial interface. HALO-based image analysis identified S100A9-positive regions of interest near the basal epithelial layer **(Fig. 6F)**. S100A9 mean intensity was similarly increased in the P-OPL stromal regions close to the basal epithelium **(Fig. 6F, G)**. These findings support the activation of S100A9-associated myeloid/inflammatory programs and epithelial interferon response features in P-OPL lesions.

**Fig. 6.**
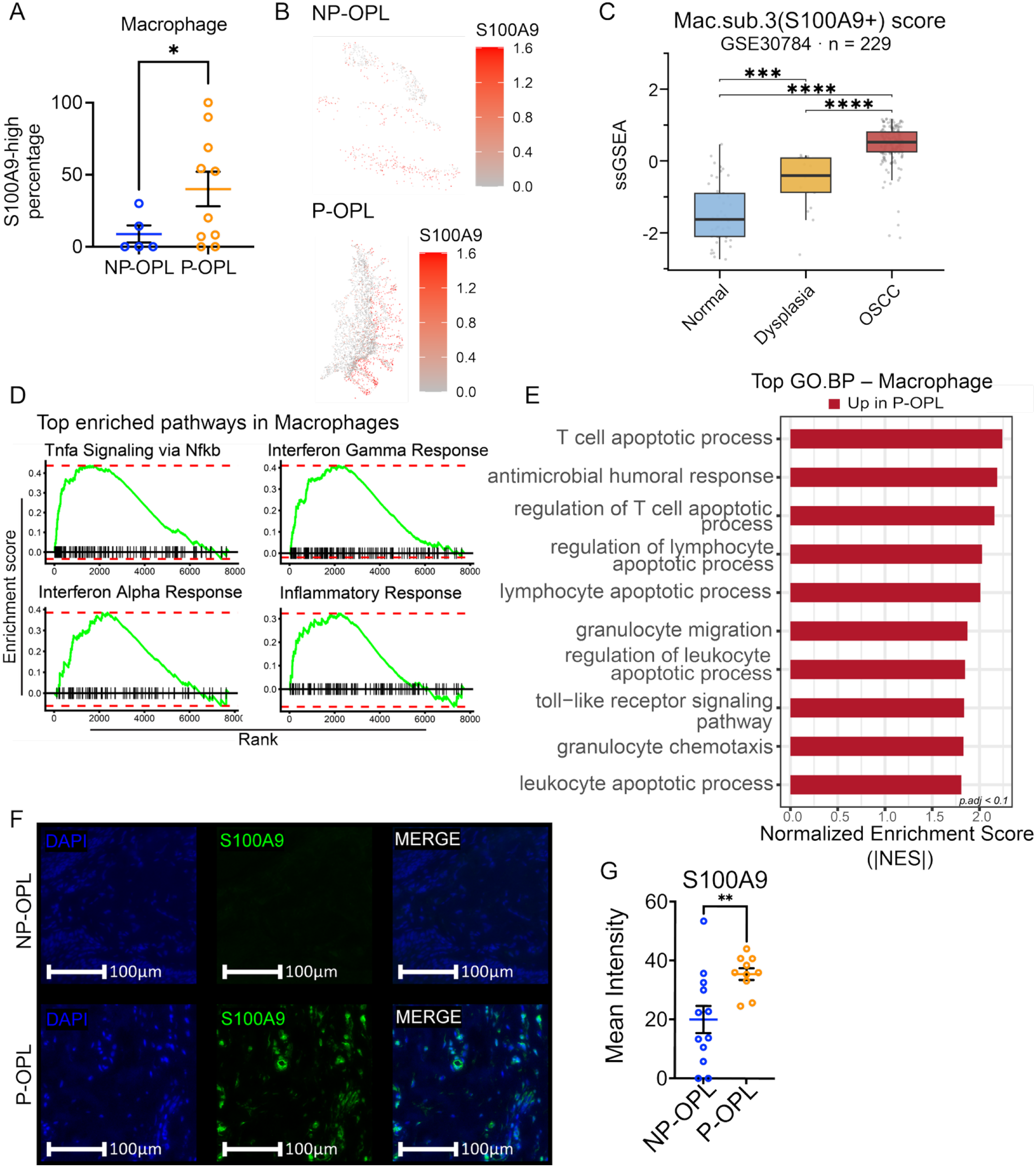
Validation of S100A9-associated macrophage programs in P-OPL. **A** S100A9-high macrophages were defined as cells with normalized *S100A9* expression greater than the cohort-wide median value. Percentage of S100A9-high macrophages was calculated as the number of S100A9-high macrophages divided by the total number of macrophages in that biospecimen. Each point represents one independent biospecimen. **B** Representative Xenium feature plots showing *S100A9* expression in NP-OPL and P-OPL samples. **C** Boxplots of ssGSEA scores of Mac.sub.3 markers in normal, dysplasia, and OSCC samples showing elevation along with malignant progression (GSE30784; n = 229; 45 normal/ 17 dysplasia/ 167 OSCC). Groups were compared by two-tailed Mann–Whitney tests across all three pairwise contrasts with Benjamini–Hochberg correction. \*\*\**q* < 0.001, \*\*\*\**q* < 0.0001. **D** GSEA of MSigDB Hallmark gene sets in macrophages (based on pseudobulk DESeq2 results). **E** GSEA of Gene Ontology Biological Process gene sets in macrophages (based pseudobulk DESeq2 results). **F** Representative immunofluorescence images of S100A9 staining in epithelial and stromal compartments near the epithelial interface. Scale bars, 100 μm. **G** Quantification of stromal S100A9 mean intensity (n= 12 for NP-OPL and n=10 for P-OPL). The *p* value for panel A was calculated based on GEE model. **(A, G)** P-OPL and NP-OPL were compared by two-tailed Mann–Whitney test. \**p* < 0.05, \*\**p* < 0.01, \*\*\**p* < 0.001, \*\*\*\**p* < 0.0001. Data are presented as mean ± SEM.

#### T-cell validation

Next, we assessed whether the TIGIT-associated T-cell program identified in the spatial analysis was reproduced in the independent scRNA-seq cohort (n=12; four NP-OPLs, eight P-OPLs, as T cells were detected only in these samples in the cohort). The frequency of TIGIT-high T cells and the mean exhaustion-associated transcriptional score were both increased in P-OPL compared to NP-OPL (p<0.05) **(Fig. 7A, B; Fig. S17C)**. 17 exhaustion-associated markers, including *PDCD1, CTLA4, HAVCR2, LAG3* and *TIGIT*, were generally elevated in the P-OPL T cells **(Fig. 7C)**. The exhaustion scores also increased across the overall T-cell population and within T-cell subsets, including mucosa-associated invariant T (MAIT) cells, γδ T cells (gd T), CD4 T cells, CD8 T cells, and Tregs **(Fig. 7D, E)**. Validation in the external cohort (GSE30784; n = 229; 45 normal/ 17 dysplasia/ 167 OSCC) showed a specific enrichment of ssGSEA score of the T cell exhaustion markers in OSCC samples, while an increasing trend was observed from normal to dysplasia samples **(Fig. 7F).** Broadly, 11 of 17 T cell exhaustion markers were confidently detected (mean intensity above the 25th percentile and variance above the 10th percentile of the array-wide distribution). Relatively to normal samples, *HAVCR2, TIGIT,* and *ENTPD1* exhibited significant association with progression, while *CTLA4, NR4A1, TOX, CD38,* and *CXCL13* were particularly enriched in OSCC samples **(Fig. 7G)**. Among these T cell exhaustion markers, *TOX* showed specific upregulation in OSCC samples, and *TIGIT* showed tight progression association, with stage-wise elevation from normal to dysplasia and thereafter to OSCC samples **(Fig. 7H)**. GSEA based on the pseudobulk results further showed enrichment of KRAS signaling and interferon-gamma response programs in P-OPL T cells **(Fig. S20; Table S30-31)**. Together, these findings validate the enrichment of TIGIT-associated exhaustion programs in P-OPL T-cells.

**Fig. 7.**
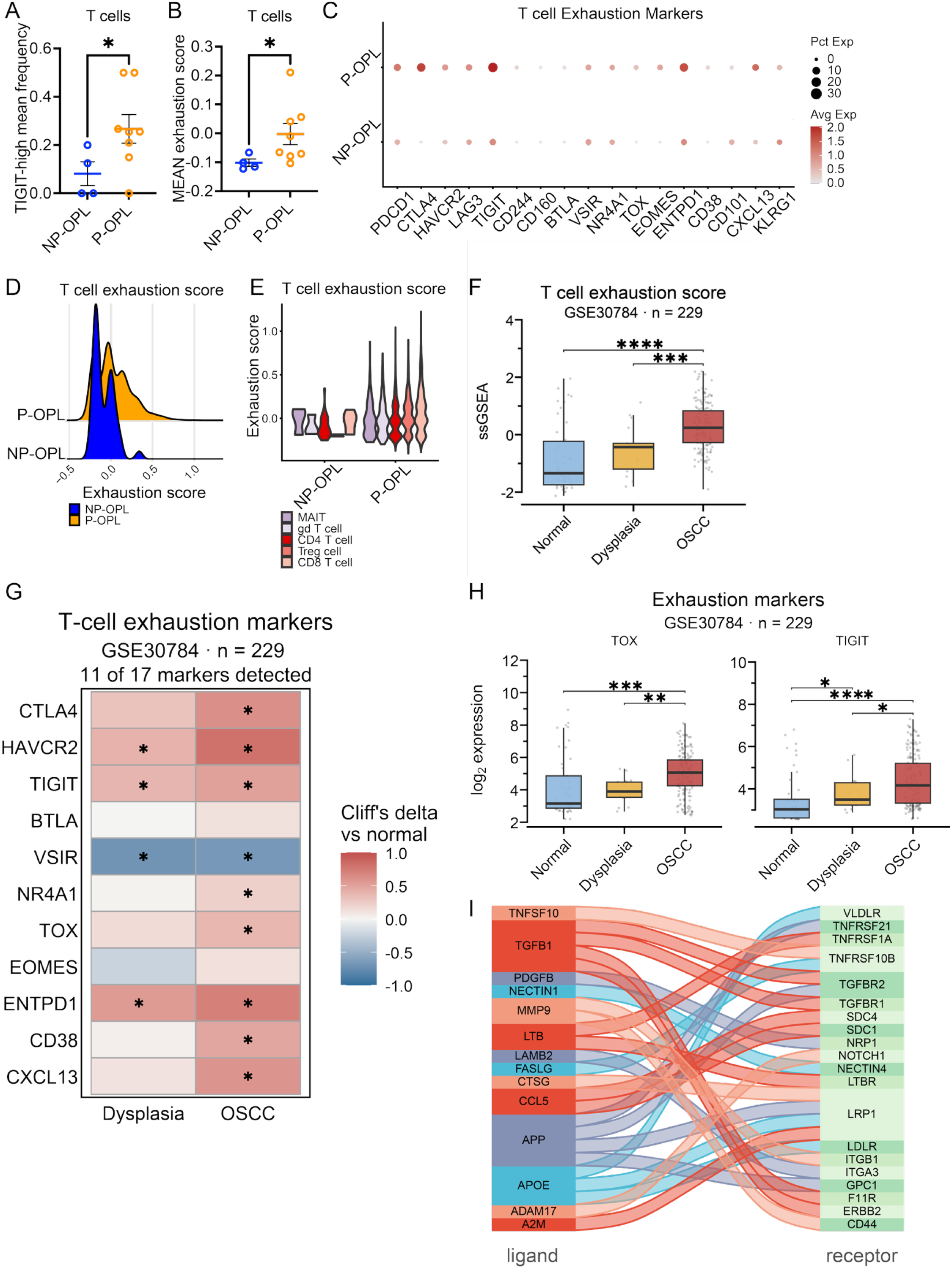
T cells in P-OPL show increased TIGIT expression and exhaustion-associated programs. **A** Mean frequency of TIGIT-high T cells in the independent whole-transcriptome scRNA-seq validation cohort. TIGIT-high cells were defined as T cells in the top quartile of *TIGIT* expression. P-OPL samples showing increased TIGIT-high T cell frequency compared with NP-OPL samples. **B** Mean T-cell exhaustion score in NP-OPL and P-OPL samples. Exhaustion scores were computed using Seurat AddModuleScore with a gene exhaustion signature. **C** Dot plot showing expression of individual T cell exhaustion markers in NP-OPL and P-OPL T cells. Dot size represents the percentage of expressing cells, and color represents average expression. **D** Ridgeline plot showing the distribution of T cell exhaustion scores across NP-OPL and P-OPL T cells. **E** T-cell exhaustion scores across T cell subtypes, including MAIT cells, γδ T cells, CD4 T cells, regulatory T cells, and CD8 T cells. **F** Boxplots of Single-sample Gene Set Enrichment Analysis (ssGSEA) scores of T cell exhaustion markers in normal, dysplasia, and OSCC samples showing scores increase along with malignant progression (GSE30784; n = 229; 45 normal/ 17 dysplasia/ 167 OSCC). **G** Heatmap of individual T cell exhaustion markers in dysplasia, and OSCC samples showing effect size (Cliff’s delta) of each gene in dysplasia or OSCC relative to normal (GSE30784; n = 229; 45 normal/ 17 dysplasia/ 167 OSCC). **H** Boxplots of log2 expression of *TOX* and *TIGIT* in normal, dysplasia, and OSCC samples showing an increase in expression along with malignant progression (GSE30784; n = 229; 45 normal/ 17 dysplasia/ 167 OSCC). **I** NicheNet alluvial diagram showing prioritized immune-to-epithelial ligand–receptor interactions predicted in P-OPL compared with NP-OPL. Immune populations were used as ligand-producing sender cells, and epithelial cells were used as receptor-expressing receiver cells. **(A–E)** Data were analyzed in the independent scRNA-seq validation cohort restricted to biospecimens in which T cells were recovered (n = 12; 4 NP-OPL and 8 P-OPL); three biospecimens (1 NP-OPL, 2 P-OPL) yielded no T cells. **I** was analyzed in the full single-cell RNA-seq validation cohort (n = 15; 5 NP-OPL and 10 P-OPL). **(A**, **B)** Statistical significance was assessed using two-tailed Mann–Whitney tests. Data are presented as mean ± SEM. \**p* < 0.05, \*\**p* < 0.01, \*\*\**p* < 0.001, \*\*\*\**p* < 0.0001. **(F–H)** Groups were compared by two-tailed Mann–Whitney tests across all pairwise contrasts with Benjamini–Hochberg correction. \**q* < 0.1, \*\**q* < 0.01, \*\*\**q* < 0.001, \*\*\*\**q* < 0.0001.

#### Dendritic-cells validation

We performed a pseudobulk analysis on DCs in P-OPL vs. NP-OPL in the scRNA-seq validation cohort (n=15; 5 NP-OPL, 10 P-OPL) **(Table S32)**. Pathway analysis of differentially expressed genes revealed enrichment of the interferon-α/γ response, TNF-α/NF-κB signaling, cholesterol homeostasis, and inflammatory response programs in P-OPL **(Fig. S21; Table S33).**

Collectively, these validation analyses demonstrated that the progression-associated programs identified by Xenium spatial transcriptomics are reproducible across independent FFPE single-cell and immunofluorescence cohorts. Rather than reflecting an isolated marker change, P-OPL lesions showed a coordinated epithelial– immune remodeling program across the basal epithelial, myeloid, dendritic cell, and T-cell compartments. This program included inflammatory and stress-associated activation of KRT14⁺/KRT15⁺ basal epithelial cells marked by *MX1*, *NOTCH3*, *IGFBP7*, *CD44*, and *CXCL14*; S100A9-linked myeloid remodeling with TNF-α/NF-κB, interferon, toll-like receptor, granulocyte migration, and inflammatory response programs; enrichment of TIGIT-high and exhaustion-associated T-cell states; and inflammatory activation of dendritic cells together with altered positioning relative to the epithelial interface. Together, these findings support a model in which lesions that progress to OSCC have already undergone epithelial–immune niche reprogramming before overt malignant transformation, characterized by basal epithelial interferon/stress signaling, S100A9-associated myeloid inflammation, checkpoint-associated T-cell dysfunction, and disruption of dendritic cell immune surveillance.

### Immune-to-epithelial ligand-receptor signaling highlights inflammatory and remodeling niches in P-OPL

To investigate immune-to-epithelial signaling during progression, we applied NicheNet using immune cell populations as ligand-producing sender cells and epithelial cells as the receptor-expressing receiver population and compared P-OPL with NP-OPL (**Fig. 7I; Fig. S22; Table S34**). As epithelial cells were modeled as the receiver population, we first examined the expression of cognate receptors in epithelial cells. The epithelial receiver cells expressed receptor programs involved in TGF-β signaling, TNF/death-receptor signaling, EGFR/ERBB signaling, integrin-mediated adhesion, Notch/nectin signaling, and LRP-family lipid/scavenger receptor pathways, including *TGFBR1/2*, *TNFRSF* family members, *EGFR*, *ERBB2/3*, *ITGA/ITGB* family members, *NOTCH1*, *NECTIN1/4*, *CD44*, *LRP1*, *LDLR*, and *VLDLR* **(Fig. S22A)**.

We next examined whether prioritized immune-derived ligands predicted to regulate epithelial transcriptional programs were increased in P-OPL. The candidate ligands included *TGFB1*, *TNFSF10*, *FASLG*, *CCL5*, *NECTIN1*, *LAMB2*, *MMP9*, *APOE*, *ANXA1*, *CXCL9*, *CXCL13*, *CXCL16*, *TIMP1*, *GPNMB*, and *ICAM1* **(Fig. S22B)**. These ligands are distributed across multiple immune-sender populations. Myeloid populations contributed to remodeling and immune-regulatory ligands, including *TGFB1*, *MMP9*, *TIMP1*, *APOE*, *ANXA1*, *GPNMB*, and *NECTIN1*, whereas lymphoid populations contributed to inflammatory and cytotoxic ligands, including *CCL5*, *TNFSF10*, and *FASLG* **(Fig. S22B)**.

The predicted ligand–receptor interaction map highlighted several signaling modules consistent with epithelial stress, immune regulation, and tissue remodeling. These included TGFB1–TGFBR/integrin-associated signaling, TNFSF10/FASLG–TNFRSF/FAS death-receptor signaling, CCL5/CXCL9/CXCL13/CXCL16-associated inflammatory chemokine signaling, MMP9/TIMP1/ADAM/LAMB2-associated extracellular matrix and adhesion signaling, NECTIN1/Notch-associated epithelial interaction pathways, and APOE–LRP family receptor signaling **(Fig. 7I; Fig. S22C; Table S34)**.

Together, NicheNet analysis identified a predicted immune-to-epithelial communication network in P-OPL in which myeloid and lymphoid sender populations converged on epithelial receptor programs associated with inflammatory signaling, death-receptor signaling, chemokine activity, extracellular matrix remodeling, adhesion, nectin/Notch signaling, and lipid/scavenger receptor pathways. These predicted interactions overlapped with the independently observed spatial and transcriptional features of P-OPL, including increased TIGIT-associated T-cell states, S100A9-linked epithelial and myeloid remodeling, altered dendritic-cell organization, and epithelial– immune niche restructuring. Thus, immune-to-epithelial ligand–receptor inference nominates candidate signaling axes that may contribute to the inflammation and remodeling of the epithelial niche observed in progressive lesions.

### Bootstrap-validated LASSO selection yields a stable, parsimonious model for progression risk

The expanded clinical analysis evaluated six candidate predictors and retained mild versus non-mild histologic grade as the sole predictor. The clinical model achieved an apparent AUC of 0.682 and bootstrap optimism-corrected AUC of 0.648. The expanded clinical plus biomarker analysis evaluated 14 candidate predictors, comprising six clinical variables and all eight available MX1, S100A9, and TIGIT readouts. In the regularized regression model fitted to the complete 23-patient cohort, the selected elastic-net mixing parameter was 0.1, corresponding to an *L*_2_-dominant penalty that allowed correlated biomarker readouts to retain non-zero coefficients simultaneously. Full-pipeline bootstrap stability analysis identified a recurrent core comprising histologic grade, S100A9 positive-cytoplasmic intensity, and mean count of TIGIT^+^ T cells in 200 μm of the epithelial dysplastic interface, which were selected in 76.6%, 70.8%, and 67.5% of 1,000 bootstrap resamples, respectively. The expanded clinical plus biomarker model achieved an apparent AUC of 0.917 and bootstrap optimism-corrected AUC of 0.808. These findings identify histologic grade, S100A9 cytoplasmic intensity, and mean count of TIGIT^+^ T cells in 200 μm of the epithelial dysplastic interface as the recurrent core of a progression-associated clinical plus biomarker signature that improves the clinical-only model AUC.

In a separate, hypothesis-driven clinical plus spatial analysis using the prespecified reduced candidate set, one-standard-error LASSO selected histologic grade, log1p-transformed dendritic cell density, and CD3-positive T-cell average distance from the epithelial interface as predictors. These predictors were subsequently refitted using a ridge-stabilized logistic regression, yielding an apparent AUC of 0.894 and a bootstrap optimism-corrected AUC of 0.839. These findings suggest that dendritic-cell abundance and epithelial T-cell spatial organization provide complementary progression-associated information beyond the histologic grade.

## DISCUSSION

This study identified spatial epithelial–immune niche remodeling as an early feature of oral malignant transformation. Outcomes in OSCC remain strongly dependent on the stage at diagnosis, and risk assessment for oral premalignant lesions continues to rely heavily on histologic grading despite known limitations in reproducibility and predictive performance (*1, 12, 13, 37, 38*). Prior genomic and single-cell studies have defined the mutational, epithelial, stromal, and immune complexities of established HNSCC (*3, 39, 40*). However, the spatial organization of epithelial–immune interactions that distinguishes progressive from non-progressive premalignant lesions before invasion remains poorly resolved. By integrating FFPE-compatible Xenium spatial transcriptomics with independent FFPE single-cell RNA sequencing and immunofluorescence validation, our study extends this literature to the premalignant window and identifies coordinated changes in the epithelial state, immune-cell function, and tissue geometry that distinguish progressive from non-progressive OPL.

A central finding was the emergence of stress-, interferon-, and remodeling-associated basal epithelial programs in at-risk lesions. The epithelial subcluster enriched for *MX1*, *CDKN2B*, *NOTCH3*, *PTGS1*, and *NELL2* expanded in P-OPL and OSCC, consistent with interferon-responsive epithelial activation, NOTCH3-associated epithelial plasticity, and prostaglandin-linked inflammatory signaling. Independent scRNA-seq validation further supported the enrichment of related basal epithelial programs involving *MX1*, *NOTCH3*, *IGFBP7*, *CD44*, and *CXCL14* in the KRT14⁺ and KRT15⁺ epithelial compartments. These findings suggest that basal epithelial cells in progressive lesions are not passive targets of immune remodeling but active participants in shaping an inflammatory and immunoregulatory premalignant niche. Conversely, the epithelial program marked by *CEACAM6*/*CEACAM8*/*CEACAM1* and *TMPRSS2* was increased in P-OPL and OSCC. Previous studies have linked CEACAM-family molecules to epithelial mucosal signaling, mucosal immune regulation, epithelial– neutrophil biology, and EGFR-dependent OSCC invasion, whereas TMPRSS2 has been detected in oral epithelial tissues and embedded in IL-13-associated secretory epithelial programs in the airway epithelium (*23, 24, 41, 42*).

Myeloid remodeling is the second major axis of progression. The macrophage subcluster marked by *APOE*, *MMP12*, *S100A9*, *ISG15*, and *SDC1* was enriched in OSCC and reflected inflammatory, interferon-responsive, matrix remodeling, and immunoregulatory programs. Previous studies have supported the role of S100A9 and S100A8/9 heterodimer (calprotectin) in myeloid-derived immune suppression, inflammatory tumor promotion, and innate immune signaling, and implicated ISG15 in tumor-associated macrophage polarization and immune modulation (*26–29*). In the independent scRNA-seq cohort, P-OPL lesions showed increased odds of S100A9-high macrophage enrichment, and pathway analyses supported the activation of TNF-α/NF-κB, interferon, toll-like receptor, granulocyte migration, and inflammatory response programs. These findings support a model in which S100A9-associated myeloid inflammation is already established in progressive premalignant lesions, rather than appearing only after invasive cancer develops.

T-cell remodeling adds a checkpoint-associated dimension to the progressive niche. *TIGIT* increased across the NP-OPL to P-OPL to the OSCC continuum in the Xenium data, and the independent scRNA-seq cohort reproduced increased TIGIT-high T-cell frequency and higher exhaustion-associated transcriptional scores. This aligns with previous studies showing that T-cell exhaustion and immune checkpoint expression can emerge during oral carcinogenesis and oral premalignancy, as well as OSCC-specific studies implicating TIGIT in intratumoral lymphocyte biology and broader work defining the TIGIT/CD226 axis in antitumor immunity (*10, 11, 43, 44*). Together with increased epithelial–immune separation, these data suggest that checkpoint-associated T-cell dysfunction may emerge before frank invasion and contribute to ineffective immune surveillance during oral malignant transformation. These transcriptional and checkpoint-expression findings are consistent with an exhaustion-associated state but do not demonstrate functional T-cell exhaustion.

Importantly, the S100A9 axis was supported by functional assays. S100A9 and the combined recombinant S100A8 and S100A9 promoted wound closure, DNA synthesis, and proliferation in oral epithelial and OSCC models, whereas inhibition of TLR4 and RAGE attenuated these effects. These findings are consistent with the known biology of S100A8 and S100A9 as the myeloid alarmin complex calprotectin and with prior evidence that S100A8 and S100A9 can signal through TLR4- and RAGE-associated pathways (*35, 36, 45*). These experiments provided a functional bridge between the S100A9-enriched inflammatory microenvironment observed in P-OPL and epithelial behaviors relevant to malignant transformation, including wound closure, proliferation, and tissue remodeling. Thus, S100A9 is not only a marker of inflammatory myeloid remodeling in our dataset but also a candidate mediator of epithelial remodeling through receptor-dependent signaling.

Dendritic cell dynamics suggest that immune activation and constraints coexist during progression. We observed an increased representation of a *CD1A*/*CD74*/*FCER1A*/*S100B*/*RUNX3*-positive tissue-resident DC-like subset of cells in P-OPL and OSCC, alongside contraction of a *NOTCH1*/*CD1C*-associated cDC2-like program. Because conventional human CD1C⁺ dendritic cells are linked to antigen presentation and T-cell priming, this pattern may reflect reduced productive epithelial immune surveillance during progression (*46, 47*). These changes suggest that antigen-sampling or inflammatory DC states may persist, while productive T-cell priming contracts, and immune cells become more spatially separated from the basal epithelium as lesions advance.

Spatial geometry has emerged as a measurable feature of the early immune exclusion. In P-OPL and OSCC, T cells and dendritic cells were positioned farther from the epithelial basal layer than in NP-OPL, and nearest-neighbor analysis showed enrichment of TIGIT-high T cells and loss of dendritic cells within the epithelial neighborhoods. These spatial metrics capture a core attribute of immune escape that cannot be inferred from the cell abundance alone. They suggested that the risk of progression in OPL may be encoded not only by the presence of epithelial and immune cells but also by where they are positioned relative to the basal epithelial compartment. NicheNet analysis adds a mechanistic layer to the spatial and transcriptional changes observed during progression by suggesting that P-OPLs are characterized by coordinated immune-epithelial signaling rather than isolated alterations in individual cell populations. Myeloid-derived ligands, including *TGFB1*, *MMP9*, *TIMP1*, *APOE*, *ANXA1*, *GPNMB*, and *NECTIN1*, and lymphoid-derived ligands, including *CCL5*, *TNFSF10*, and *FASLG*, were predicted to converge on epithelial receptor programs associated with TGF-β signaling, death-receptor activity, inflammatory chemokine responses, integrin-mediated adhesion, extracellular matrix remodeling, nectin/Notch signaling, and LRP-family lipid and scavenger receptor pathways. This convergence provides a plausible mechanistic bridge between independently observed S100A9-associated myeloid remodeling, TIGIT-associated T-cell dysfunction, altered dendritic cell organization, and progressive separation of immune cells from the epithelial interface. These findings suggest that immune cells in progressive lesions may not simply become less abundant or more spatially excluded, but may also actively reshape epithelial stress, survival, adhesion, and remodeling programs through multiple partially overlapping signaling inputs.

The inferred network further identified testable signaling axes—including TGFB1–TGFBR, TNFSF10/FASLG– death-receptor, MMP9/TIMP1–matrix-remodeling, NECTIN1/Notch-associated, ANXA1-associated, and APOE-LRP-family pathways, for mechanistic investigation and potential interception. However, NicheNet infers communication from coordinated ligands, receptors, and target gene expression; it does not establish direct ligand–receptor engagement, confirm the assignment of sender and receiver populations, or demonstrate causality. Therefore, these interactions should be considered as hypotheses for subsequent validation using spatial colocalization, protein-level analysis, and functional perturbation.

Together, these findings support a working model in which progressive OPLs acquire a checkpoint-enriched, myeloid-skewed, spatially excluded epithelial–immune niche before overt malignant transformation. This niche is characterized by a NOTCH3/IFN/prostaglandin-associated basal epithelial program, S100A9-associated myeloid remodeling, contraction of a NOTCH1/CD1C-associated cDC2-like program, TIGIT-associated T-cell dysfunction, and measurable increases in epithelial–immune distance. This model shifts the interpretation of OPL progression from a purely epithelial dysplasia process to a spatially organized tissue-ecosystem transition involving both molecular state and microanatomic organization.

To strengthen generalizability, we validated the key findings across independent modalities. The FFPE scRNA-seq validation cohort reproduced progression-associated epithelial shifts involving *MX1*, *NOTCH3*, *IGFBP7*, *CD44*, and *CXCL14*, S100A9-linked myeloid remodeling, and TIGIT-associated T cell exhaustion. Immunofluorescence validation further supported increased *MX1*, *S100A9*, and TIGIT-associated immune features, and altered DC positioning in P-OPL. In the independent bulk microarray cohort spanning normal mucosa, dysplasia, and OSCC (GSE30784, n = 229; 45 normal/ 17 dysplasia/ 167 OSCC), the same epithelial, myeloid, and T-cell exhaustion signatures increased along the disease axis. These cross-platform findings support the conclusion that progressive oral premalignant lesions harbor a coordinated epithelial–immune remodeling program before overt malignant transformation.

The translational implication is that spatial epithelial immune features may complement conventional histologic grading. In our proof-of-concept biomarker model, the addition of S100A9 cytoplasmic intensity and %TIGIT to histologic grade improved apparent and optimism-corrected discrimination compared with the clinical-only model. Although this model requires larger prospective validation, it illustrates how molecular and spatial immune features can be integrated with histology to identify high-risk lesions before overt morphological transformation. Candidate biomarkers include epithelial *NOTCH3*/*MX1* and immune *TIGIT*/*S100A9*, together with automated spatial features, such as epithelial–immune distance and k-nearest-neighbor neighborhood composition. Therapeutically, our data raise testable concepts to rebalance checkpoint-associated T-cell dysfunction, restrain S100A9-linked myeloid remodeling, and restore dendritic cell/T-cell communication at the epithelial interface.

This study has several limitations. The Xenium discovery dataset uses a targeted immune-oncology panel rather than whole-transcriptome spatial profiling, which may miss additional epithelial, stromal, vascular, metabolic, or neuronal programs. Cohort size was modest, particularly for subgroup analyses by sex, age, lesion site, exposure history, histologic grade, and clinical metadata was incomplete for some samples. Ligand–receptor analyses are inferential and require functional validation. The classifier was exploratory, was developed in only 23 patients, and was evaluated solely by internal resampling; it has not undergone independent prospective validation. Future studies should evaluate these spatial biomarkers in larger longitudinal OPL cohorts with standardized sampling, prospective outcome annotation, and orthogonal validation using multiplex imaging or clinical-grade assays. Functional perturbation experiments are also important to test whether the inferred S100A9, TIGIT, Notch, and matrix-remodeling axes directly contribute to epithelial remodeling, immune exclusion, or malignant transformation.

In conclusion, by integrating high-plex Xenium spatial transcriptomics with FFPE single-cell RNA sequencing, immunofluorescence validation, ligand–receptor inference, functional S100A9 perturbation, and exploratory biomarker modeling, we resolved progression-associated epithelial and immune states together with their microanatomy. Progressive lesions showed expansion of a NOTCH3/IFN/prostaglandin epithelial program, enrichment of TIGIT-high T cells and S100A9-high macrophages, contraction of a NOTCH1/CD1C-associated cDC2-like program, altered dendritic cell positioning, and increased epithelial–immune distances. These convergent molecular and geometric readouts suggest practical biomarkers and actionable pathways for biomarker-guided surveillance and precision interception in oral cancer.

## Acknowledgments

The authors thank the JP Sulzberger Columbia Genome Center and the University of Michigan Advanced Genomics Core for their assistance. Library preparation and next-generation sequencing were performed in the BRCF Advanced Genomics Core at the University of Michigan. Single-cell RNA sequencing reported in this publication was supported by the National Cancer Institute of the National Institutes of Health under Award Number P30CA046592 through the use of the University of Michigan Single Cell and Spatial Analysis Shared Resource. Xenium spatial transcriptomics was performed using the Genome Engineering Core (GECO) at the JP Sulzberger Columbia Genome Center, Columbia University Irving Medical Center.

## Author contributions

Conceptualization: YJ, FMH; Methodology: YJ, ODK, FMH; Investigation: YJ, IA, FMH; Data curation: YJ, FMH; Formal analysis: YJ, IA, FMH; Visualization: YJ, IA, FMH; Resources: EP, JMG, FMH; Writing— original draft: YJ, FMH; Writing—review and editing: YJ, IA, JMG, EP, ODK, FMH; Supervision: FMH.

## Data, code, and materials availability

All data needed to evaluate and reproduce the results in the paper will be presented in the paper and the Supplementary Materials. The whole transcriptome scRNA-seq and spatial transcriptomics (Xenium) datasets generated in this study will be deposited in the NCBI Gene Expression Omnibus (GEO) and accession numbers will be provided upon acceptance.

